# Combining GWAS and TWAS to identify candidate causal genes for tocochromanol levels in maize grain

**DOI:** 10.1101/2022.04.01.486706

**Authors:** Di Wu, Xiaowei Li, Ryokei Tanaka, Joshua C. Wood, Laura E. Tibbs-Cortes, Maria Magallanes-Lundback, Nolan Bornowski, John P. Hamilton, Brieanne Vaillancourt, Christine H. Diepenbrock, Xianran Li, Nicholas T. Deason, Gregory R. Schoenbaum, Jianming Yu, C. Robin Buell, Dean DellaPenna, Michael A. Gore

**Affiliations:** Plant Breeding and Genetics Section, School of Integrative Plant Science, Cornell University, Ithaca, NY 14853, USA; Department of Crop & Soil Sciences, Institute of Plant Breeding, Genetics, & Genomics, University of Georgia, Athens, GA 30602, USA; Department of Agronomy, Iowa State University, Ames, IA 50011, USA; Department of Biochemistry and Molecular Biology, Michigan State University, East Lansing, MI 48824, USA; Department of Plant Biology, Michigan State University, East Lansing, MI 48824, USA; Department of Plant Sciences, University of California, Davis, CA 95616, USA; United States Department of Agriculture, Agricultural Research Service, Wheat Health, Genetics, and Quality Research Unit, Pullman, WA 99164, USA

## Abstract

Tocochromanols (tocopherols and tocotrienols, collectively vitamin E) are lipid-soluble antioxidants important for both plant fitness and human health. The main dietary sources of vitamin E are seed oils that often accumulate high levels of tocopherol isoforms with lower vitamin E activity. The tocochromanol biosynthetic pathway is conserved across plant species but an integrated view of the genes and mechanisms underlying natural variation of tocochromanol levels in seed of most cereal crops remains limited. To address this issue, we utilized the high mapping resolution of the maize Ames panel of ∼1,500 inbred lines scored with 12.2 million single-nucleotide polymorphisms to generate metabolomic (mature grain tocochromanols) and transcriptomic (developing grain) data sets for genetic mapping. By combining results from genome- and transcriptome-wide association studies, we identified a total of 13 candidate causal gene loci, including five that had not been previously associated with maize grain tocochromanols: four biosynthetic genes (*arodeH2* paralog, *dxs1*, *vte5*, and *vte7*) and a plastid S-adenosyl methionine transporter (*samt1*). Expression quantitative trait locus (eQTL) mapping of these 13 gene loci revealed that they are predominantly regulated by *cis*-eQTL. Through a joint statistical analysis, we implicated *cis*-acting variants as responsible for co-localized eQTL and GWAS association signals. Our multi-omics approach provided increased statistical power and mapping resolution to enable a detailed characterization of the genetic and regulatory architecture underlying tocochromanol accumulation in maize grain and provided insights for ongoing biofortification efforts to breed and/or engineer vitamin E and antioxidant levels in maize and other cereals.

## Introduction

Tocochromanols, which include the biosynthetically related tocopherols and tocotrienols, are a group of plant-synthesized lipid-soluble antioxidants that have a chromanol ring derived from homogentisic acid (HGA) and isoprenoid-derived hydrophobic side chains. The saturated side chain of tocopherols is derived from phytyl-diphosphate (PDP), whereas the tocotrienol side chain has three double bonds and is derived from geranylgeranyl diphosphate (GGDP). Tocopherols and tocotrienols occur as four biosynthetically related isoforms (α, β, δ, and γ) that vary in the degree and position of methyl groups on their chromanol rings. Among the tocochromanols, α-tocopherol has the highest vitamin E activity (DellaPenna and Mène-Saffrané 2011), while tocotrienols tend to have greater antioxidant activity (Sen *et al*. 2006). Although severe vitamin E deficiency leading to ataxia and myopathy is rare in human populations (Traber 2012), suboptimal dietary vitamin E intake exists in certain population segments (Ford *et al*. 2006; McBurney *et al*. 2015) and has been linked to an elevated risk of cardiovascular diseases (Knekt *et al*. 1994; Kushi *et al*. 1996). Tocochromanols are found at high levels in plant seeds where they confer protection against lipid peroxidation during seed storage and germination (Sattler *et al*. 2004). However, α-tocopherol is not the major tocochromanol in most cereal seed oils, which limits the dietary vitamin E intake of both humans and animals (DellaPenna and Mène-Saffrané 2011).

Tocochromanols are only synthesized by photosynthetic organisms, and the tocochromanol biosynthetic pathway is highly conserved in the plant kingdom (DellaPenna and Mène-Saffrané 2011). In the committed step of tocopherol synthesis (Figure 1), a homogentisate phytyltransferase (VTE2) condenses PDP and HGA from the shikimate pathway to produce 2-methyl-6-phytyl-1,4-benzoquinol (MPBQ) (Sattler *et al*. 2004). In the monocot lineage, HGA can also be condensed with geranylgeranyl diphosphate (GGDP) by homogentisate geranylgeranyltransferase (HGGT1) to generate 2-methyl-6-geranylgeranyl-1,4-benzoquinol (MGGBQ), the committed step for tocotrienol synthesis. MPBQ and MGGBQ are substrates for a series of methylations by MPBQ/MGGBQ methyltransferase (VTE3) and γ-tocopherol methyltransferase (VTE4) and cyclization by tocopherol cyclase (VTE1), whose sequence and numbers of reactions generate the α, β, δ, and γ isoforms of tocopherols and tocotrienols (Shintani and DellaPenna 1998; Porfirova *et al*. 2002; Van Eenennaam *et al*. 2003; Cheng *et al*. 2003; Sattler *et al*. 2004). While GGDP for tocotrienol synthesis comes directly from the isoprenoid pathway, the generation of PDP for tocopherol synthesis is more complex and still not completely resolved. Though differences in leaf tocopherol synthesis exist between monocots and dicots, tocopherol synthesized in seed requires chlorophyll biosynthesis (Diepenbrock *et al*. 2017; Zhan *et al*. 2019) and the activity of VTE7, an alpha/beta hydrolase that interfaces with chlorophyll synthesis to release phytol (Albert *et al*. in review). Phytol is then sequentially phosphorylated to PDP through the action of phytol kinase (VTE5) and phytol phosphate kinase (VTE6) (Valentin *et al*. 2006; Vom Dorp *et al*. 2015).

**Figure 1.**
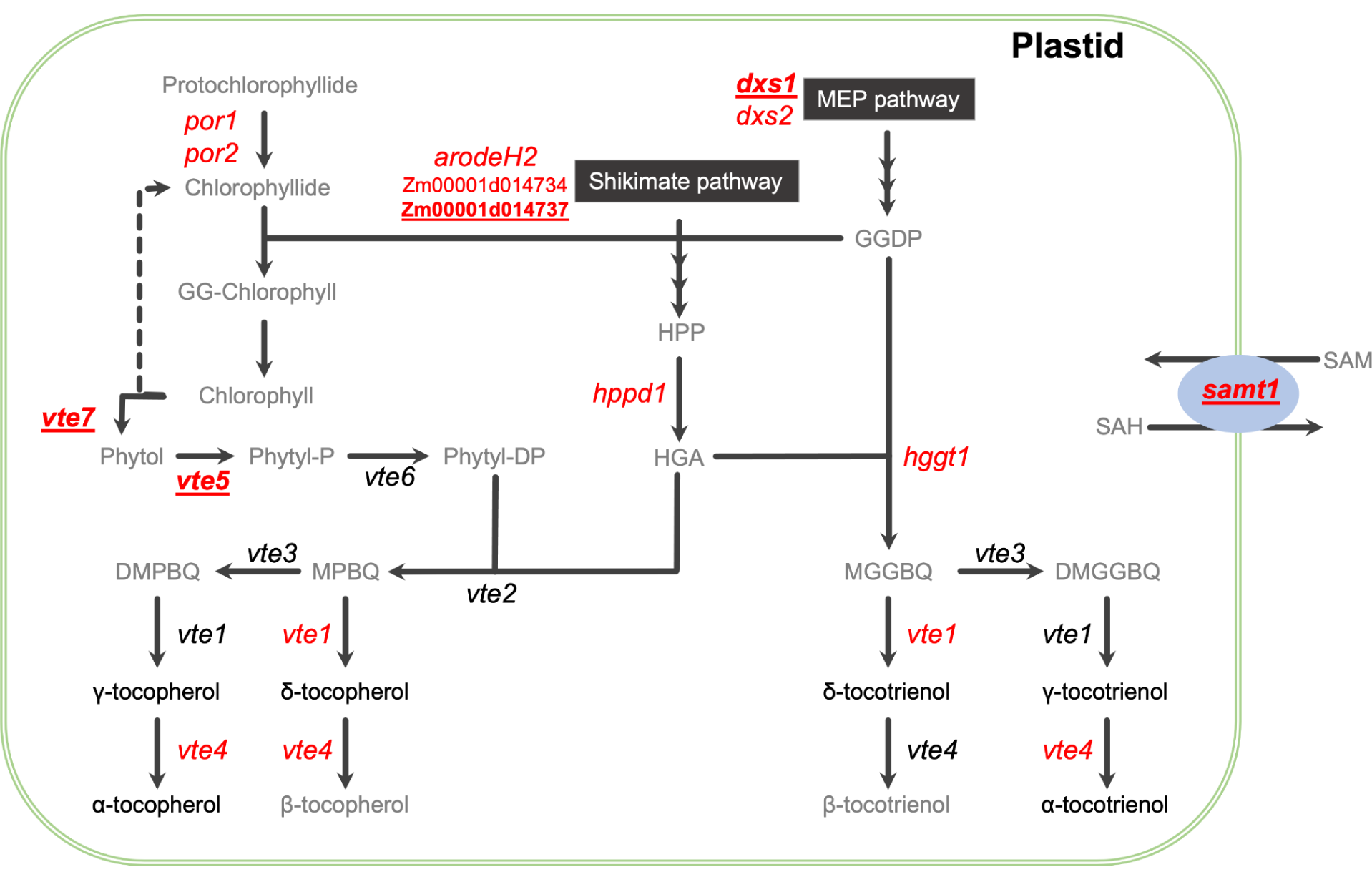
Tocochromanol biosynthetic pathways in maize. The two precursor pathways are represented as black boxes. The six quantified compounds are indicated in black non-italicized text. The names of key *a priori* genes are italicized at the pathway step(s) catalyzed by their encoded enzyme, with the 13 candidate causal gene loci identified in this study in red text. The five genes not previously shown to be associated with tocochromanols in maize grain are red, bolded, and underlined. Compound abbreviations: DMGGBQ, 2,3-dimethyl-5-geranylgeranyl-1,4-benzoquinol; DMPBQ, 2,3-dimethyl-6-phytyl-1,4-benzoquinol; GGDP, geranylgeranyl diphosphate; GG-Chlorophyll, geranylgeranyl-chlorophyll a; HGA, homogentisic acid; HPP, p-hydroxyphenylpyruvate; MEP, methylerythritol 4-phosphate; MGGBQ, 2-methyl-6-geranylgeranyl-1,4-benzoquinol; MPBQ, 2-methyl-6-phytyl-1,4-benzoquinol; Phytyl-DP, phytyl diphosphate; Phytyl-P, phytyl monophosphate; SAM, S-adenosyl-L-methionine; SAH, S-adenosyl-L-homocysteine. Gene abbreviations: 1-deoxy-D-xylulose-5-phosphate synthase (*dxs1* and *2*); α-/β-hydrolase family protein (*vte7*); arogenate/prephenate dehydrogenase family protein (*arodeH2*); phytol kinase (*vte5*); phytol phosphate kinase (*vte6*); *p*-hydroxyphenylpyruvate dioxygenase (*hppd1*); protochlorophyllide reductase (*por1* and *2*); homogentisate geranylgeranyltransferase (*hggt1*); homogentisate phytyltransferase (*vte2*); MPBQ/MGGBQ methyltransferase (*vte3*); γ-tocopherol methyltransferase (*vte4*); S-adenosylmethionine transporter 1 (*samt1*); tocopherol cyclase (*vte1*). The *vte7* locus consists of tandemly duplicated genes (Zm00001d006778 and Zm00001d006779) in B73 RefGen_v4.

In the past decade, several loci associated with natural variation for the content and composition of tocochromanols in maize grain have been identified via genome-wide association studies (GWAS) in mapping panels. Several studies have reported strong associations between *vte4* and α-tocopherol concentration in maize grain (Li *et al*. 2012; Lipka *et al*. 2013; Wang *et al*. 2018; Baseggio *et al*. 2019), with relatively weaker associations detected for *vte1*, *hggt1*, and an arogenate/prephenate dehydratase with maize grain tocotrienol levels (Lipka *et al*. 2013; Baseggio *et al*. 2019). Wang *et al*. (2018) implicated genes outside of the core tocochromanol biosynthetic pathway as playing a role in maize grain tocopherol levels, including genes involved in fatty acid biosynthesis, chlorophyll metabolism, and chloroplast function. While these combined studies provided some insight into the genetic basis of tocochromanol levels in maize grain, these studies were limited by mapping panel size and marker density.

Through a joint-linkage (JL) analysis and GWAS in the 5,000-line U.S. maize nested association mapping (NAM) panel with high statistical power, Diepenbrock *et al*. (2017) identified 50 unique QTL for tocochromanol grain traits. Of these, 13 QTL were resolved to seven *a priori* pathway genes (*dxs2*, *sds*, *arodeH2*, *hppd1*, *hggt1*, *vte3*, and *vte4*) and six non-pathway genes (*por1*, *por2*, *snare*, *ltp*, *phd*, and *fbn*) that encoded predicted functions not previously associated with tocochromanol traits. Although maize grain is a non-green, non-photosynthetic tissue, two protochlorophyllide reductases (POR1 and POR2) were found to be major loci controlling total tocopherols and were hypothesized to be part of a cycle that provides chlorophyll-derived phytol for tocopherol synthesis in the maize embryo, which contain exceedingly low, but detectable, levels of chlorophylls. The involvement of *por2* in tocopherol accumulation in maize grain was transgenically confirmed by overexpression (Zhan *et al*. 2019). Although the U.S. NAM panel provided unparalleled mapping resolution for these 13 QTL, 37 QTL for tocochromanols with moderate effects or limited contrasting genotypes in the NAM panel could not be resolved to the gene level, thus other mapping approaches combined with more diverse mapping panels could have substantial utility in identifying additional underlying candidate causal genes.

For terminal phenotypes such as grain tocochromanols, intermediate phenotypes or endophenotypes can offer orthogonal genetic information to better connect genotype to phenotype. In a gene-based association approach that can generate insights into biological processes, a transcriptome-wide association study (TWAS) correlates mRNA abundance with complex trait variation, which allows for the statistical connection of an intermediate phenotype to a terminal phenotype (Hirsch *et al*. 2014; Pasaniuc and Price 2017; Lin *et al*. 2017; Li *et al*. 2021). In an assessment of TWAS for the genetic dissection of tocochromanol and carotenoid grain traits in the Goodman-Buckler association panel, Kremling *et al*. (2019) showed that the statistical power to detect previously identified causal genes could be increased through an ensemble approach combining GWAS and TWAS results with the Fisher’s combined test (FCT). Additionally, this approach was used to identify plausible causal genes associated with natural variation for water use efficiency traits in sorghum (Pignon *et al*. 2021; Ferguson *et al*. 2021). The genetic markers used in GWAS could also be linked to mRNA abundance via expression quantitative trait locus (eQTL) mapping (Majewski and Pastinen 2011), enabling the regulatory landscape of traits to be better explored as has been conducted for oil content and composition in maize grain (Li *et al*. 2013).

In this study, we conducted a comprehensive genetic dissection of tocochromanol grain phenotypes in a large maize association panel (∼1,500 inbreds) that leveraged the scoring of ∼12.2 million SNP markers and transcript abundances of more than 22,000 genes from developing grain. Our integrated GWAS and TWAS approach combined with eQTL mapping to pinpoint multiple candidate causal genes and uncover their regulatory control.

## Materials and methods

### Experimental design for genetic mapping

A collection of 1,815 maize inbred lines from the Ames panel (Romay *et al*. 2013) was grown as a single replicate at Iowa State University in Ames, IA, in 2015 and 2017. The Ames panel was arranged in an augmented complete block design. For each year, two blocking directions were assigned: each range block consisted of three adjacent rows of plots, and each pass block consisted of eight adjacent columns of plots. At least one B73 check plot was planted within each pass and range block. The inbred lines were grouped into two and three tiers for 2015 and 2017, respectively, according to their days to silking (flowering time) recorded in Romay *et al*. (2013) for the 2015 design and pollination date in 2015 for the 2017 design. Experimental units were one-row 3.05 m plots having ∼18 plants, with 0.76 m inter-row spacing and a 0.76 m alley. Approximately six plants per plot were self-pollinated and hand-harvested at physiological maturity. The kernels from all dried and shelled ears were bulked to form a representative sample. For each plot, 25 kernels were ground in an IKA Tube Mill Control (IKA-Werke, Staufen, Germany) and ground tissue stored in cryovials at -80 °C.

### Phenotypic data analysis

The extraction of tocochromanols from 15-20 mg of ground kernels and their quantification by high-performance liquid chromatography (HPLC) and fluorometry were as previously described (Lipka *et al*. 2013). Briefly, tocopherols and tocotrienols were assessed on 3,539 grain samples from 1,762 inbred lines and the repeated B73 check plots. The nine evaluated tocopherol and tocotrienol phenotypes in μg g^−1^ dry seed were as follows: α-tocopherol (αT), δ-tocopherol (δT), γ-tocopherol (γT), α-tocotrienol (αT3), δ-tocotrienol (δT3), γ-tocotrienol (γT3), total tocopherols (ΣT, calculated as αT + δT + γT), total tocotrienols (ΣT3, calculated as αT3 + δT3 + γT3), and total tocochromanols (ΣTT3, calculated as ΣT + ΣT3). Statistical outliers were identified and filtered from the raw HPLC data, followed by a mixed linear model analysis that modeled genetic and non-genetic (field and laboratory) effects to produce best linear unbiased estimator (BLUE) values for the 1,762 inbred lines (Supplementary Table S1) and heritability estimates as described in the Supplementary Methods. Considering that morphologically extreme grain types can potentially have inflated tocochromanol concentrations based on a dry sample weight basis, we conservatively excluded 265 inbred lines that had been classified as sweet corn, popcorn, or having other endosperm mutations.

### Genotypic data

We used the genotype data processing and BEAGLE v5.0 (Browning *et al*. 2018) imputation approaches implemented by Wu *et al*. (2021) as described in the Supplementary Methods. The generated imputed marker data set for enabling GWAS of 1,462 inbred lines with both genotypic and phenotypic data consisted of 14,613,169 SNPs from maize HapMap 3.2.1 (Bukowski *et al*. 2018) in B73 RefGen_v4 coordinates. These SNP loci were further filtered for quality to produce a high quality set of 12,184,805 SNPs that had minor allele frequency (MAF) ≥ 1% and predicted dosage *r^2^* (DR2) ≥ 0.80 (Supplementary Data Set 1) for performing GWAS with the mixed linear model. In PLINK version 1.9 (Purcell *et al*. 2007) with a sliding window of 100 kb and step size of 25 SNPs, the complete set of 12,184,805 SNPs was LD pruned to construct two reduced marker sets (Supplementary Data Set 1): 1) 7,319,895 SNPs with pairwise *r^2^* < 0.99 for performing GWAS with a multi-locus mixed model (MLMM), and 2) 344,469 SNPs with pairwise *r*^2^ < 0.10 for estimation of population structure and relatedness.

### Genome-wide association study

We conducted GWAS of the nine tocochromanol phenotypes scored on the 1,462 lines with a previously described procedure (Wu *et al*. 2021). In brief, to correct for heteroscedasticity and non-normality of error terms, the Box-Cox power transformation procedure (Box and Cox 1964) was used to select an optimal value of convenient lambda for transforming the non-negative BLUE values of each phenotype (Supplementary Table S2). Given that several negative BLUE values were generated in the model fitting process, we added a constant that made all values positive and no less than 1E−09 before applying the transformation (Supplementary Table S2). Each of 12,184,805 SNPs was tested for an association with transformed BLUE values from the 1,462 lines (Supplementary Table S3) using a mixed linear model (Yu *et al*. 2006) that employed the population parameters previously determined approximation (Zhang *et al*. 2010) in the R package GAPIT version 2018.08.18 (Lipka *et al*. 2012). In GAPIT, the reduced set of 344,469 SNPs was used to calculate the kinship matrix with the VanRaden method I (VanRaden 2008) and principal components (PCs). The optimal number of PCs to include in the mixed linear model fitted for each phenotype was determined by the Bayesian information criterion (BIC) ^(^Schwarz 1978). The likelihood-ratio-based R^2^ statistic (R^2^_LR_) (Sun *et al*. 2010) was used to approximate the amount of phenotypic variation explained by a SNP. The “*p.adjust*” function in base R version 4.0.2 (R Core Team 2018) was used to apply the false discovery rate (FDR) multiple test correction procedure (Benjamini and Hochberg 1995) to the *P*-values of tested SNPs.

To control for large-effect loci, we used the MLMM approach of Segura *et al*. (2012) as implemented by Wu *et al*. (2021) to conduct a GWAS of each transformed tocochromanol phenotype with the reduced set of 7,319,895 SNPs that alleviated model constraints by removing perfectly correlated SNPs. In each model, we included the same kinship matrix and BIC-determined optimal number of PCs that had been used for GWAS with the mixed linear model.

### Experimental design for transcriptomic profiling

In 2018, 1,023 of the 1,815 maize inbred lines from the Ames panel evaluated for grain tocochromanols plus five additional founders of the U.S. maize NAM panel (Yu *et al*. 2008; McMullen *et al*. 2009) were grown as a single replicate at Iowa State University. This germplasm set was initially constructed by including 256 lines that met at least one of the following criteria: 1) extreme (high or low) for a grain metabolite phenotype in the 2015 field trial; 2) founder of the U.S. NAM panel; or 3) available genome assembly. The additionally randomly selected 771 lines were included to increase genetic diversity and sample size. The 1,028 non-check lines were partitioned and randomized into 24 augmented incomplete blocks based on pollination dates recorded in the 2015 and 2017 field trials and divided across two tiers. A B73 check plot was planted within each block to control for spatial variation across the field. Additionally, two local checks were planted at random positions in each block to account for temporal variation across fresh harvest dates that spanned more than a month. Within each block, the line that had flowered the latest in previous years was selected to serve as one of the two local checks; ties were broken by choosing the line with the highest sample call rate based on a filtered partially imputed GBS data set (Supplementary Data Set 2). Each selected local check was also planted in their adjacent later-flowering block, so that two local check lines were present in blocks 2-24. An additional early-flowering line (S 117) was identified as a local check and planted in block 1, ensuring that two local check lines were also planted in block 1. In addition, the 25 local checks (S 117, C38, A508, A641 Goodman-Buckler, C31, 807, LH202, 764, PHG71, PHB47, SD101, A680, B93, NC292, NC280, LH208, H100, NC252, NC314, LH51, CI 187-2, NC324, Mo11, NC334, and M37W) were planted in a separate third tier to account for field effects on these lines. Experimental units were one-row plots of the same dimensions and plant numbers as used in the genetic mapping experiments. Of the ∼six pollinated ears per plot, a single self-pollinated ear was hand-harvested at ∼23 days after pollination (DAP), followed by immediately freezing the dehusked ear in liquid nitrogen and keeping it covered in dry ice (and, for some samples, frozen at -80°C) until shelling. The 23 DAP time point was selected to capture maximal increase in tocochromanols and strong expression of known pathway genes (Diepenbrock *et al*. 2017). To control for temporal effects, a self-pollinated ear of a local check from the third tier was hand-harvested at ∼23 DAP on each day of fresh harvest, with all harvested ears identically processed with liquid nitrogen and dry ice prior to shelling. The mid-section of each frozen ear was individually shelled on dry ice and its kernels stored at −80°C. In total, 1,012 non-check and 107 check kernel samples were collected.

### RNA isolation and 3**′** mRNA Sequencing

Eight to ten frozen kernels per sample were ground using liquid nitrogen cooled grinding cups in an IKA Tube Mill Control (IKA-Werke, Staufen, Germany) and ∼100 mg of ground tissue was used for RNA isolation using a modified hot borate method (Wan and Wilkins 1994). RNA samples were DNase treated and checked for quality per Hershberger *et al*. (2022). RNA samples were randomized into 96-well plates and shipped overnight on dry ice to the Genomics Facility of the Cornell Institute of Biotechnology. Included in each plate submission were positive controls consisting of the same pool of B73 control RNA aliquoted into four wells in each plate, as well as four negative controls per plate consisting of water. Libraries were constructed using the Lexogen QuantSeq 3′ mRNA-Seq Library Kit FWD (Lexogen, Greenland, NH) and sequenced on an Illumina NextSeq 500 producing 85 nt single-end reads (Illumina, San Diego, CA) with each plate being split in half and each half being sequenced on a single lane to achieve desirable coverage.

### Expression abundance determination

The 3′ QuantSeq reads were cleaned using two rounds of Cutadapt version 2.3 (Martin 2011) to trim Illumina adapters, the first 12 bases, and polyA tails. Reads were then aligned to the B73 RefGen_v4 reference genome (Jiao *et al*. 2017) using HISAT2 version 2.1.0 (Kim *et al*. 2019) with the following parameters: --min-intronlen 20, --max-intronlen 60000, --dta-cufflinks, and -- rna-strandness F. The resultant alignments were then sorted using SAMTools version 1.9 (Li *et al*. 2009). Counts were then generated using the htseq-count function within HTSeq version 0.11.2 (Anders *et al*. 2015) using the B73 version 4.59 annotation with the following parameters: --format=bam, --order=pos, --stranded=yes, --minaqual=10, --idattr=ID, --type=gene, and -- mode=union. The DESeq2 *rlog* function (Love *et al*. 2014) was used to normalize the count data for the set of 1,171 RNA samples (1,119 kernel samples plus 52 positive controls). All genes with a normalized count of less than or equal to zero in all samples were removed from the final count matrix. Several stringent quality control measures based on sampling concerns (e.g., moldy kernels etc.), alignment rate, between sample correlation value, genotype confirmation assessment, and heterozygosity level were implemented to filter out low quality samples as described in the Supplementary Methods and summarized in Supplementary Table S4, resulting in a final set of 741 high-quality samples that were used for subsequent analysis.

### Expression data analysis

The expression data set consisting of 665 samples for 664 non-check lines and 76 samples for 25 check lines was further stringently filtered for statistical outliers at the gene level following the approach of Hershberger *et al*. (2022) to ensure high quality data for statistical analysis. The filtering steps and metrics are summarized in Supplementary Table S4. With the filtered expression data set, we fit a mixed linear model that enabled the modeling of genetic and non-genetic effects as described in the Supplementary Methods. Of the 664 lines, we excluded 104 classified as sweet corn, popcorn, or having other endosperm mutations and an additional 15 lines not analyzed in GWAS. The final data set contained BLUE expression values of 22,136 genes across 545 lines.

To account for inferred confounders that influence expression variation, the PEER approach (Stegle *et al*. 2012) was separately applied to the 545 line × 22,136 gene matrix of BLUE expression values as previously described (Hershberger *et al*. 2022). The contribution of 11 PEER hidden factors was subtracted to generate a residual data set of the BLUE expression values (hereafter, PEER values). Studentized deleted residuals (Neter *et al*. 1996) were used to identify and remove significant outliers (Bonferroni α = 0.05) from the set of PEER values (Supplementary Data Set 3).

### Transcriptome-wide association study

We conducted TWAS on the 545 lines and 22,136 genes with a mixed linear model approach (Yu *et al*. 2006; Zhang *et al*. 2010). Briefly, a mixed linear model was fit for the combination of each tocochromanol phenotype (transformed BLUE values, response variable) and expressed gene (outlier-screened PEER values, explanatory variable) using the “*GWAS*” function and “P3D” option set to FALSE in the R package rrBLUP version 4.6 (Endelman 2011). To construct the SNP marker set for the 545 lines, 12,018,644 biallelic SNPs (DR2 ≥ 0.80; MAF ≥ 1%) were subsetted from the full set of 14,613,169 SNPs and pruned down to 328,892 SNPs with pairwise *r^2^* < 0.10 with a 100 kb sliding window and 25 SNP step size in PLINK version 1.9 (Supplementary Data Set 4). The PCs and the kinship matrix were generated from the 328,892 SNPs as described above. The optimal models for all tocochromanol phenotypes included kinship and no PCs, as determined by the BIC (Schwarz 1978). The TWAS for the *vte7* locus was conducted separately, given that reads for the *vte7* locus were uniquely processed to account for tandemly duplicated genes (Supplementary Methods).

### Fisher’s combined test

The top 10% of the smallest *P*-value SNPs (1,218,480 SNPs) from GWAS with the mixed linear model were selected to perform FCT following the method of Kremling *et al*. (2019). In brief, the GWAS *P*-value of each top 10% SNP was assigned to its nearest gene based on the B73 RefGen_v4 assembly and B73 v4.59 annotation and then paired with the TWAS *P*-value for that gene. For genes not tested in TWAS, their TWAS *P*-values were set to 1 before combining with GWAS *P*-values. For each gene, FCT was conducted with the “*sumlog*” function implemented in the R package metap version 1.1 (Dewey 2019).

### Candidate gene identification

Given that GWAS, TWAS, and FCT differ in their statistical power and type of independent variables, we did not directly compare *P*-value thresholds across methods but instead used the rankings of *P*-values following that of Kremling *et al*. (2019). The top 0.02% of SNPs were selected according to their *P*-value from GWAS results for each phenotype, with selection of the percentage threshold guided by the oligogenic genetic architecture of these phenotypes in the U.S. maize NAM panel (Diepenbrock *et al*. 2017). Considering the rapid LD decay in the Ames panel (Romay *et al*. 2013), candidate genes were identified within 100 kb of the peak SNP for each locus following the method of Wu *et al*. (2021). The top 0.5% of genes according to their *P*-value were selected from TWAS and FCT results for each phenotype, resulting in a total number of unique genes identified across phenotypes by each method comparable to that of GWAS.

The identification of candidate genes was assisted by a list of 126 *a priori* candidate genes involved in the accumulation of grain tocochromanol levels (Supplementary Table S5). The physical positions of 50 unique JL-QTL common support intervals (CSIs) and GWAS markers associated with the nine tocochromanol phenotypes in the U.S. NAM panel (Diepenbrock *et al*. 2017) were uplifted via Vmatch version 2.3.0 (Kurtz 2010) to B73 RefGen_v4 coordinates (Supplementary Tables S6, S7) as described in Wu *et al*. (2021). A BLASTP with default parameters was conducted to identify the top hits of undescribed candidate causal genes (Supplementary Table S8) in Arabidopsis and rice as previously described (Wu *et al*. 2021).

### eQTL mapping

We performed expression QTL (eQTL) mapping of the identified candidate causal genes. To conduct eQTL mapping, the 12,018,644 SNPs subsetted in the TWAS approach were individually tested for association with PEER values of each candidate causal gene using a mixed linear model implemented in GAPIT version 2018.08.18 (Lipka *et al*. 2012) in R version 4.0.2 (R Core Team 2018). The calculated PCs and kinship matrix used in TWAS were used in eQTL mapping, with the optimal number of PCs determined by BIC (Schwarz 1978). To have a stringent control of the Type I error rate in the presence of complex LD patterns and strong association signals, we accounted for multiple testing with a 5% Bonferroni adjusted significance threshold (*P*-value ≤ 4.16E−09), with peak SNPs of loci identified as described in Wu *et al*. (2021).

### Annotation of variants

SNP effect analysis (SnpEff; Cingolani *et al*. 2012) was conducted following the approach of Diepenbrock *et al*. (2021) to predict the effects of GWAS-associated SNPs located within candidate causal genes. To quantitatively assess whether variant sites are evolutionarily conserved, genomic evolutionary rate profiling (GERP; Davydov *et al*. 2010) scores available from two earlier studies (Kistler *et al*. 2018; Ramstein *et al*. 2020) were extracted for the same SNP sites within candidate causal genes.

### eCAVIAR

To quantify the probability that a variant was responsible for both GWAS and *cis*-eQTL signals, we used the eQTL and GWAS CAusal Variants Identification in Associated Regions (eCAVIAR) method of Hormozdiari *et al*. (2016) that accounts for LD patterns and allelic heterogeneity when implementing a probabilistic model for integrating GWAS and *cis*-eQTL results. The eCAVIAR approach was applied to candidate causal gene loci detected via GWAS that had a significant *cis*-eQTL signal. For each gene-phenotype pair, the *t*-values of all significant GWAS and eQTL SNPs within 100 or 250 kb of the candidate causal gene and pairwise LD matrices calculated from these SNPs in PLINK 1.9 (Purcell *et al*. 2007) were used as the input data sets for the eCAVIAR software. The ± 250 kb window was only used for two genes (*arodeH2* Zm00001d014734 and *vte1*) that had a peak eQTL signal >100 kb from its respective gene. Given the potential of allelic heterogeneity, the maximum number of causal SNPs at a locus was set to three. A stringent colocalization posterior probability (CLPP) cutoff threshold of ≥ 0.01 was used to identify SNPs that were potentially causal in both the GWAS and eQTL studies.

## Results

### Phenotypic variation

We assessed the extent of quantitative variation for tocochromanol concentrations in physiologically mature grain samples harvested from two outgrowths of the maize Ames panel. The measurement of six tocochromanol compounds by HPLC showed that γT (∼55%) and γT3 (∼23%) collectively accounted for nearly 80% of ΣTT3, whereas the α and δ isoforms for both tocopherols and tocotrienols individually represented approximately 1% (δT3) to 10% (αT3) of ΣTT3 (Table 1). The tocochromanol compound with the highest vitamin E activity, αT, had the third lowest mean concentration (5.83 μg g^−1^ dry seed) and accounted for only ∼8% of ΣTT3. Within a compound class, significant pairwise correlations between BLUE values (α = 0.05) were strongest between the δ and γ isoforms for tocopherols (*r* = 0.67) and tocotrienols (*r* = 0.62), whereas the strongest correlation between compound classes was observed for αT with αT3 (*r* = 0.45). However, only weak correlations (−0.15 to 0.19) were found between all other pairs of compounds despite having a shared biosynthetic pathway (Supplementary Figure S1). As inferred from the high estimates of heritability on a line-mean basis (0.77 to 0.94, Table 1), the majority of variation for each of the six tocochromanol compounds and three sum phenotypes was attributable to genetic variation in the Ames panel (Table 1 and Supplementary Figure S2).

**Table 1.** Means, ranges, and standard deviations (Std. Dev.) of untransformed best linear unbiased estimator (BLUE) values (in μg g^-1^) for nine tocochromanol grain phenotypes evaluated in the Ames panel and estimated heritabilities on a line-mean basis and their standard errors (Std. Err.) across two years.

| Phenotype | Number of lines | BLUEs |  |  | Heritabilities |  |
| --- | --- | --- | --- | --- | --- | --- |
|  |  | Mean | Range | Std. Dev. | Estimate | Std. Err. |
| $\alpha\text{T}$ | 1452 | 5.83 | −1.79 - 41.36 | 4.59 | 0.87 | 0.006 |
| $\delta\text{T}$ | 1456 | 1.74 | −0.33 - 14.32 | 1.62 | 0.85 | 0.007 |
| $\gamma\text{T}$ | 1458 | 42.19 | −1.32 - 158.91 | 21.34 | 0.86 | 0.006 |
| $\Sigma\text{T}$ | 1460 | 49.95 | 1.79 - 174.84 | 22.65 | 0.85 | 0.007 |
| $\alpha\text{T3}$ | 1456 | 7.87 | 0.89 - 23.39 | 3.21 | 0.77 | 0.010 |
| $\delta\text{T3}$ | 1454 | 0.93 | 0.01 - 17.05 | 0.97 | 0.94 | 0.003 |
| $\gamma\text{T3}$ | 1458 | 17.60 | −1.79 - 90.39 | 11.71 | 0.93 | 0.003 |
| $\Sigma\text{T3}$ | 1458 | 26.55 | 2.62 - 111.01 | 13.32 | 0.91 | 0.004 |
| $\Sigma\text{TT3}$ | 1460 | 77.04 | 18.13 - 205.36 | 28.08 | 0.87 | 0.006 |

### Genetic analysis of grain tocochromanol levels

We integrated GWAS and TWAS results through FCT, an ensemble approach shown to have enhanced statistical power over either GWAS or TWAS alone for the detection of causal genes associated with natural variation for tocochromanol grain phenotypes in maize (Kremling *et al*. 2019). The findings from FCT (top 0.5%), GWAS (top 0.02%), and TWAS (top 0.5%) for each phenotype were integrated with the genetic mapping results of the same grain phenotypes in the

U.S. NAM panel (Table 2), with the intent to further resolve loci previously found in the NAM panel to the level of causal genes (Figure 2 and Supplementary Figure S3). Within each analysis, a total of 720 unique genes were identified in GWAS, 676 in TWAS, and 918 in FCT across the nine tocochromanol phenotypes (Supplementary Tables S9, S10, S11). Of these, 330 (GWAS), 299 (TWAS), and 646 (FCT) genes were located within NAM JL-QTL CSIs of the nine phenotypes (Diepenbrock *et al*. 2017).

**Figure 2.**
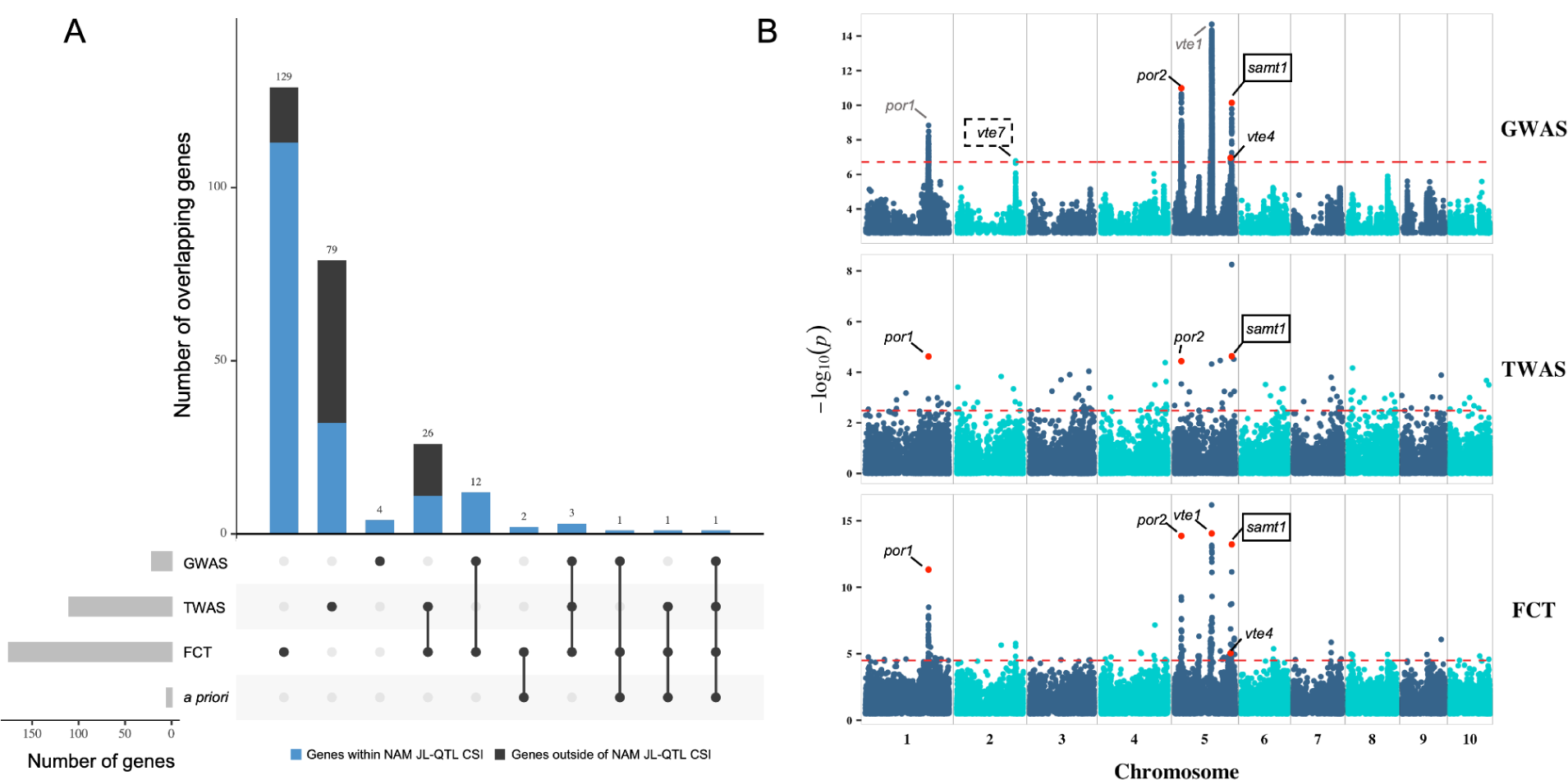
Genome-wide association study (GWAS), transcriptome-wide association study (TWAS), and Fisher’s combined test (FCT) results for δT. A, Upset plot showing the number of overlapping genes between GWAS, TWAS, FCT, and *a priori* pathway genes involved in the biosynthesis of chlorophylls and tocochromanols (Supplementary Table S5). The number of genes located within the U.S. nested association mapping (NAM) joint linkage-quantitative trait loci (JL-QTL) common support interval (CSI) for δT is highlighted in blue in the bar plots. B, Manhattan plots of GWAS, TWAS, and FCT results. Each point represents a SNP or gene with its -log_10_ *P*-value (y-axis) from GWAS, TWAS, and FCT plotted as a function of physical position (Mb, B73 RefGen_v4) across the 10 chromosomes of maize (x-axis). Red horizontal dashed lines indicate the thresholds of top 0.02%, top 0.5% and top 0.5% for GWAS, TWAS, and FCT, respectively. Causal genes (Table 2) that are within 100 kb of a top 0.02% GWAS peak SNP or ranked in the top 0.5% in TWAS or FCT are highlighted with red dots and labeled in black in the Manhattan plots. Causal genes that are within 1 Mb of a top 0.02% GWAS peak SNP are labeled in gray. Novel associations are marked with a solid line, black rectangle. Novel associations that only passed the 5% false discovery rate in GWAS are marked with a dashed line, black rectangle.

**Table 2.**
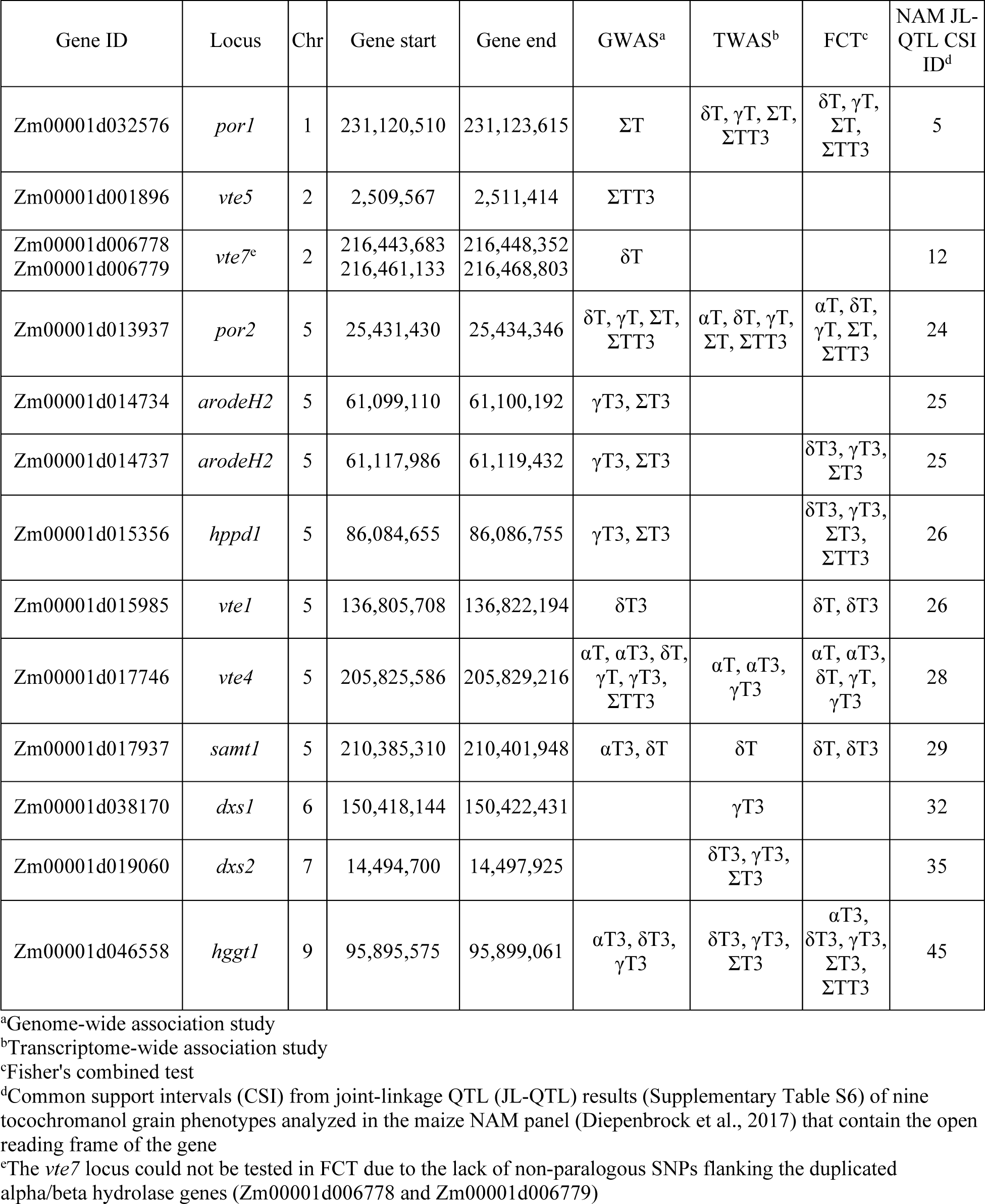
Genetic mapping results of the nine tocochromanol phenotypes in the Ames panel.

Of the 13 gene loci identified to associate with grain tocochromanols in the U.S. NAM panel by Diepenbrock *et al*. (2017), five (*por1, por2, vte4, hggt1*, and *hppd1*), which tended to be large-effect loci in the NAM panel, were detected by FCT for one or more phenotypes in the Ames panel (Table 2). Of the five genes, *por1, por2, vte4,* and *hggt1* were also identified by both GWAS and TWAS, whereas *hppd1* was only detected by GWAS. In contrast, *dxs2*, another of the 13 genes identified by Diepenbrock *et al*. (2017), was only detected by TWAS. Two copies of *arodeH2* (Zm00001d014734 and Zm00001d014737) were within 100 kb of GWAS peak SNPs for γΤ3 and ΣΤ3, with Zm00001d014734 having been previously implicated by Diepenbrock *et al*. (2017) as a gene with small allelic effects involved in the genetic control of αΤ3 and ΣΤ3. However, in the Ames panel, Zm00001d014737 was detected by both FCT and GWAS, whereas Zm00001d014734 was only detected by GWAS. In total, we reidentified seven of the 13 genes from Diepenbrock *et al*. (2017) and implicated a second *arodeH2* copy for controlling variation in tocotrienols.

The detection of these eight genes, one of which a new association, illustrated the gene-level resolution of our integrated genetic mapping approach; thus, it was applied to better resolve NAM JL-QTL CSIs and detect loci novel to the Ames panel. In total, four NAM JL-QTL CSIs were more finely dissected, resulting in novel associations with three loci (*samt1*, *vte7*, and *dxs1*) and more precise mapping of a fourth (*vte1*) not fully resolved in the U.S. NAM panel. A gene encoding a SAM transporter (*samt1*, Zm00001d017937) was detected by FCT, GWAS, and TWAS. Zm00001d017937 encodes a protein with 77% identity to Arabidopsis S-adenosylmethionine transporter 1 (SAMT1, AT4G39460) (Supplementary Table S8), which transports SAM, a tocochromanol cosubstrate for the VTE3 and VTE4 methyltransferases, through plastid envelopes and negatively impacts leaf tocopherol levels when silenced (*N*. *benthamiana*) or knocked out (Arabidopsis) (Palmieri *et al*. 2006; Bouvier *et al*. 2006). When considering the top 0.02% of SNPs most associated with δT in GWAS, the *vte7* locus consisting of tandemly duplicated genes (Zm00001d006778 and Zm00001d006779) was found to be ∼64 kb from a single associated SNP (Figure 2). This same SNP served as the peak of a δT-associated locus consisting of 45 significant SNPs at a FDR of 5% (Supplementary Table S12), providing stronger evidence for detection of *vte7* in the Ames panel compared to the NAM panel. Two additional *a priori* pathway genes, *dxs1* (TWAS) and *vte1* (FCT and GWAS), were associated with one or more tocochromanol phenotypes (Table 2). In addition to more finely dissecting NAM JL-QTL CSIs, we detected a significant association of *vte5* with ΣΤΤ3 by GWAS alone, the first report of such an association for this locus with any tocochromanol grain trait in maize. Therefore, the Ames panel not only offered gene-level resolution for existing NAM JL-QTL CSIs, but also enabled the identification of loci not detected in the NAM panel.

We also conducted a GWAS with the MLMM approach, allowing us to better resolve association signals underpinned by large-effect loci. Of the 11 gene loci detected by GWAS with the mixed linear model (Table 2), eight genes (*por2*, *vte1*, both *arodeH2* copies, *hppd1*, *vte4, samt1*, and *hggt1*) were located within 100 kb of at least one MLMM-selected SNP for one or more of the nine tocochromanol phenotypes (Supplementary Table S13). Although at slightly lower mapping resolution, the *por1* gene was located ∼162 kb from one of the multiple MLMM-selected SNPs for tocopherol phenotypes. Of the MLMM-selected SNPs within 100 kb of *vte4*, two to four SNPs each were selected for αT, αT3, and γT, whereas only a single SNP was selected for each of δT and γT3. Comparably, two to three SNPs from a ∼1.2 Mb genomic region that included *hggt1* were selected by the MLMM for δT3, γT3, and ΣT3; however, only two of the MLMM-selected SNPs were located within 100 kb of *hggt1*. As has been previously hypothesized for *vte4* in the Goodman-Buckler panel (Lipka *et al*. 2013), the selection of multiple independent SNPs by the MLMM implies that multiple causal variants (*i.e.*, allelic heterogeneity) exist at the *vte4* and *hggt1* loci in the Ames panel.

### Functional annotation of associated SNPs

We used two approaches to identify potential causal variants that change an encoded amino acid in associated genes. The SnpEff tool predicted the functional effects of the 320 GWAS-associated SNPs residing within candidate causal genes, resulting in 15 total missense variant annotations for *por1*, *por2*, *vte1*, *vte4*, *samt1*, and *hggt1* that were predicted to have moderate effects (Supplementary Table S14). Of the 15 missense variants, nine had an available GERP score from one or both of two earlier studies (Kistler *et al*. 2018; Ramstein *et al*. 2020). Each of the nine sites had at least one positive GERP score (> 0), implying as expected that these sites within coding regions are evolutionarily constrained (Davydov *et al*. 2010; Rodgers-Melnick *et al*. 2015). Collectively, six missense variants within *por1*, *vte1*, *samt1*, and *hggt1* had the highest positive GERP scores (> 2), suggesting that they are more deleterious (Yang *et al*. 2017; Lozano *et al*. 2021). Given that *vte1* was not detected by TWAS and had three of the six putatively more deleterious variants, these amino acid changes are of potentially high functional importance as they may impact the activity of tocopherol cyclase.

### eQTL mapping of candidate causal loci

To gain insights into the regulatory patterns of the loci identified through GWAS, TWAS, and FCT in the Ames panel (Table 2), eQTL mapping was conducted for each of the 13 identified candidate causal gene loci (Figure 3 and Supplementary Figure S4). Of the 13 loci, *cis*-eQTL (peak SNP within 1 Mb of gene) were identified for all but one gene (*arodeH2* Zm00001d014737), whereas a total of five *trans*-eQTL were identified for four genes (*vte5*, *por2*, *dxs1*, and *dxs2*) (Supplementary Table S15). The peak SNPs for *cis*-eQTL were within 100 kb of their respective gene, with the exceptions of *arodeH2* Zm00001d014734 (227 kb), *dxs2* (808 kb), and *vte1* (220 kb) (Supplementary Table S15). In general, *cis*-eQTL had smaller *P*-values than *trans*-eQTL, except in the case of *dxs2* where its *trans*-eQTL had a smaller *P*-value than its *cis*-eQTL. This *dxs2 trans*-eQTL was located on chromosome 6, having a peak SNP ∼1.5 kb from *phytoene synthase1* (*psy1*, Zm00001d036345), which encodes the first and committed step of carotenoid biosynthesis (Hirschberg 2001).

**Figure 3.**
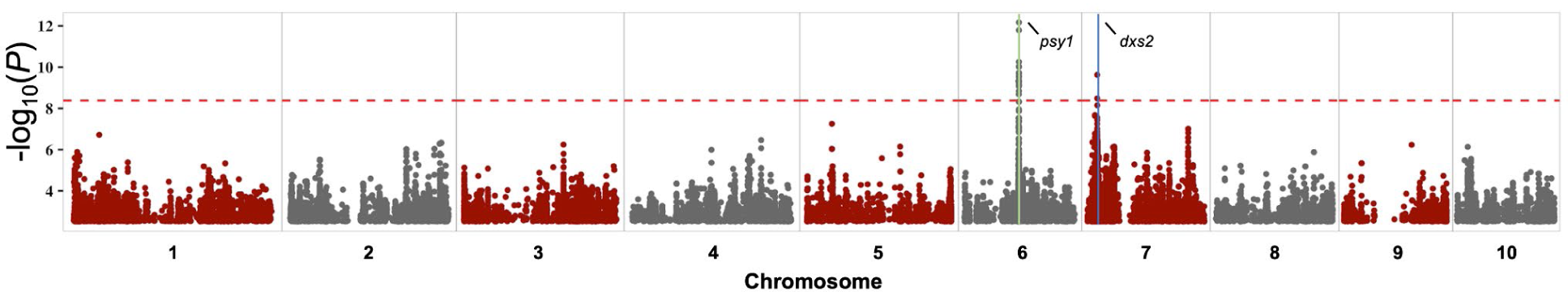
Manhattan plot of expression quantitative trait loci (eQTL) mapping results of *dxs2*. Each point represents a SNP with its -log_10_ *P*-value (y-axis) from a mixed linear model analysis plotted as a function of physical position (Mb, B73 RefGen_v4) across the 10 chromosomes of maize (x-axis). The red horizontal dashed line indicates the significant threshold after Bonferroni correction (α = 0.05). The genomic position of *dxs2* is demarcated with a blue vertical line on chromosome 7. A plausible regulating gene, *phytoene synthase1* (*psy1*), within 100 kb of the *trans*-eQTL peak SNP is indicated by a green vertical line on chromosome 6.

### Colocalization of GWAS and eQTL signals

Through the integration of GWAS and eQTL mapping results via a probabilistic approach (eCAVIAR; Hormozdiari *et al*. 2016), we tested whether a variant was responsible for both GWAS and *cis*-eQTL signals at each GWAS-identified candidate causal gene locus that had a significant *cis*-eQTL signal: *arodeH2* Zm00001d014734, *hggt1*, *hppd1*, *por1*, *por2*, *samt1*, *vte1*, *vte4*, *vte5*, and *vte7*. Of the analyzed 23 gene-phenotype pairs, 18 pairs had one to six SNPs with a colocalization posterior probability (CLPP) value—probability that a variant is causal for both a GWAS and eQTL signal—that exceeded a stringent cutoff threshold of ≥ 0.01 (Supplementary Figure S5 and Supplementary Table S16). In all, these analyses resulted in the selection of 24 unique SNPs within (*por1* and *hppd1*) or flanking (*arodeH2* Zm00001d014734, *hggt1*, *por2*, *samt1*, *vte4*, and *vte5*) eight of the 10 investigated genes. Only one of the three SNPs within a candidate causal gene was an annotated missense variant (Arg -> Gly; GERP ≥ 2), having been selected from SNPs within *por1* for total tocopherols (Supplementary Table S14). The other 21 selected SNPs were a median distance of 9.3 kb from their respective candidate causal genes.

With the strongest evidence for colocalization, a SNP (5_25434949) located 603 bp from *por2* had the highest CLPP values (0.12 - 0.17) for γT, ΣT, and ΣTT3, which was concordant given that this same SNP was the peak marker for GWAS (mixed linear model: γT, ΣT, and ΣTT3) and *cis*-eQTL signals at *por2* (Figure 4 and Supplementary Tables S9, S15). We observed additional statistical support for allelic heterogeneity (*i.e.*, independent SNPs with CLPP values ≥ 0.01) at *hggt1* for δT3 (Supplementary Figure S5), but not at *vte4* which had the same single SNP (5_205853870, ∼25 kb from *vte4*) selected for αT, αT3, δT, γT, and γT3 (CLPP values: 0.06 - 0.14). The only other GWAS detected loci that had significant *cis*-eQTL signals were *vte1* and *vte7*, but the CLPP values of SNPs for these two loci were < 0.01. Given this finding and that these two loci were not detected in TWAS, the causal variants at *vte1* and *vte7* are potentially different for the genetic control of tocochromanol and gene expression variation.

**Figure 4.**
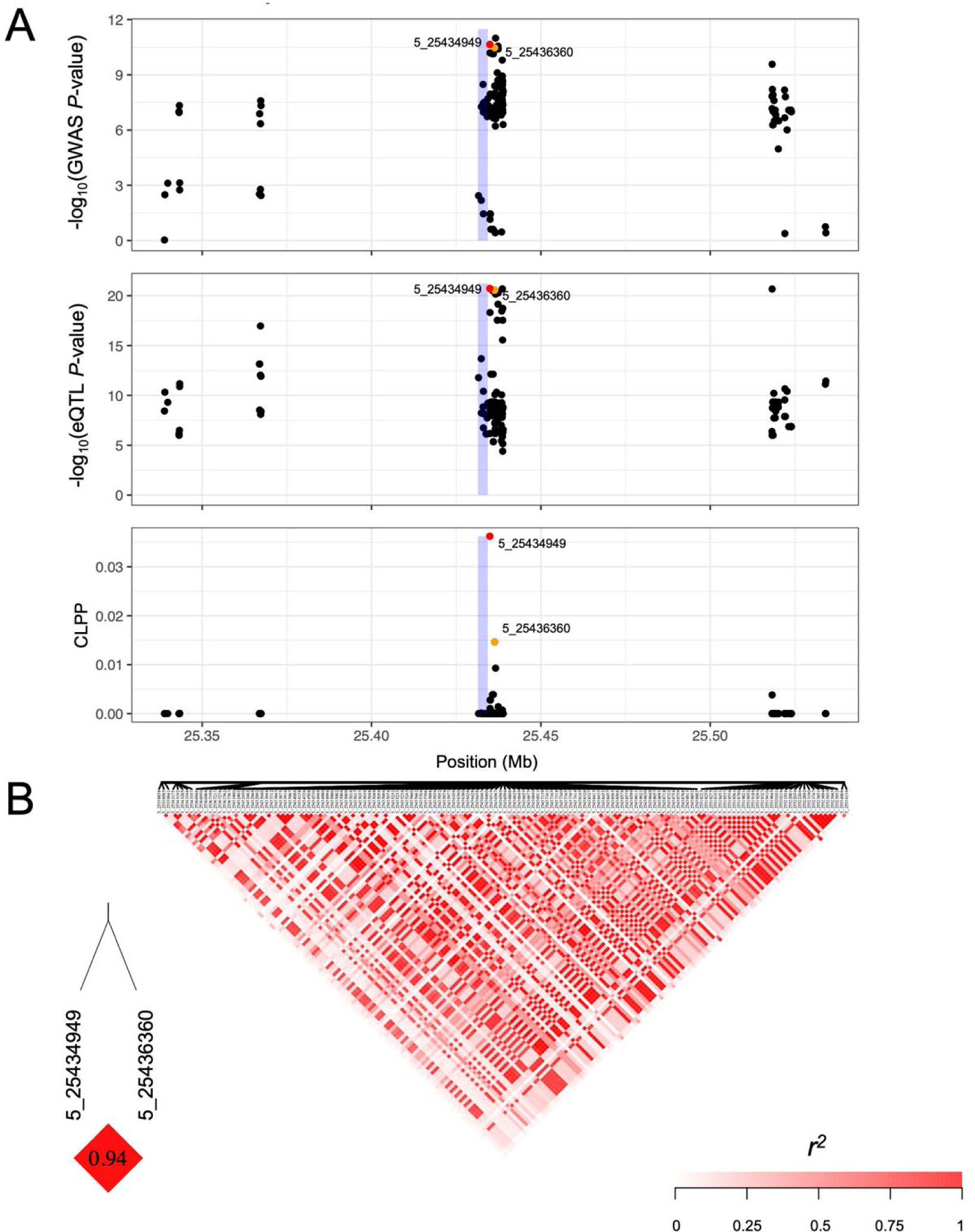
Genome-wide association study (GWAS), expression quantitative trait loci (eQTL) and eQTL and GWAS CAusal Variants Identification in Associated Regions (eCAVIAR) results at *por2* for δT. A, Three local Manhattan plots (± 100 kb) showing the GWAS, eQTL and eCAVIAR results at *por2* for δT. Each point represents a SNP with its -log_10_ *P*-value (y-axis) from a mixed linear model analysis in GWAS or eQTL, or the CLPP value from eCAVIAR, plotted as a function of physical position (Mb, B73 RefGen_v4) on chromosome 5 (x-axis). The blue rectangle represents the physical position of *por2*. The red dot represents the SNP with the highest CLPP value, and the orange dot represents the other SNP with a CLPP value passing the ≥ 0.01 threshold. B, Pairwise linkage disequilibrium (LD, *r^2^*) of all SNPs included in A, with the LD between SNPs with CLPP values ≥ 0.01 indicated in the bottom left.

## Discussion

Through combining complementary quantitative genetic approaches that span levels of biological organization, we conducted a comprehensive investigation in the maize Ames panel of nearly 1,500 lines to elucidate the genetic basis of natural variation for tocochromanol levels in grain. Imputing the Ames panel with ∼12 million SNP markers, generating transcript abundances of ∼22,000 genes at a biologically informative stage of kernel development, and HPLC profiling of more than 3,500 mature grain samples empowered implementation of GWAS, TWAS, FCT, and eQTL approaches. This enabled identification of 13 candidate causal gene loci with encoded functions responsible for varying the content and composition of grain tocochromanols. These 13 genes include a SAM transporter (*samt1*) and genes involved in synthesis of the tails of tocochromanols (*por1*, *por2*, *vte5*, *vte7*, *dxs1*, and *dxs2*), the aromatic head group (two *arodeH2* copies and *hppd1*), and the core tocochromanol pathway (*vte1*, *vte4*, and *hggt1*). Of the 13 identified genes, eight were detected by two or more of the three (GWAS, TWAS, and FCT) genetic mapping approaches (Table 2). Implicating the importance of regulatory variation, seven of the 13 genes were detected by TWAS of which two, *dxs1* and *dxs2*, were only detected by TWAS. Taken together, the integrated GWAS and TWAS method enhanced our efforts to genetically dissect grain tocochromanol phenotypes at gene-level resolution in the Ames panel.

Indicative of the high resolution and statistical power of our multi-omic approach applied to the Ames panel that contains high allelic diversity and rapid LD decay (Romay *et al*. 2013), six of the 13 genes (*samt1*, *arodeH2* Zm00001d014737, *dxs1*, *vte1*, *vte5*, and *vte7*) were not identified in the maize NAM panel (Diepenbrock *et al*. 2017). Notably, all but one of the six genes (*vte1*) had not been previously associated with maize grain tocochromanols (Li *et al*. 2012; Lipka *et al*. 2013; Diepenbrock *et al*. 2017; Wang *et al*. 2018). The association signal at *vte5* was novel to the Ames panel, likely due to a limitation of the U.S. NAM panel in its sampling of allelic diversity from only 26 diverse founder lines. In the U.S. NAM panel, *samt1*, *dxs1*, and *vte7* were each located within an unresolved NAM JL-QTL CSI, but *arodeH2* Zm00001d014737, which was detected by GWAS and FCT, was in a NAM JL-QTL CSI that was instead associated with *arodeH2* Zm00001d014734 (Diepenbrock *et al*. 2017). *vte1* was present in a NAM JL-QTL CSI that also included *hppd1*, but unlike *hppd1* the *vte1* locus could not be conclusively identified due to the recombination-suppressed pericentromeric region in which it resides, which limited mapping precision (Diepenbrock *et al*. 2017). The Ames panel, however, provided the increased resolution necessary to resolve both *hppd1* and *vte1* at the gene level. Six other smaller effect loci identified for grain tocochromanols in the U.S. NAM panel (*sds*, *vte3*, *snare*, *ltp*, *phd*, and *fbn*) were not identified in the Ames panel, suggesting that though it has superior physical resolution it has lower statistical power than a NAM design for identifying small effect QTL.

This study identified the first metabolite transporter associated with grain tocochromanols, Zm00001d017937, which encodes the maize ortholog of Arabidopsis *SAMT1*, which transports SAM into plastids (Palmieri *et al*. 2006; Bouvier *et al*. 2006). Maize *samt1* was associated with δΤ, δΤ3, and αΤ3 (Table 2), consistent with the two plastid-localized tocopherol methyltransferases (VTE3 and VTE4) being dependent on it for import of their co-substrate, SAM, from the cytosol (DellaPenna 2005; Bouvier *et al*. 2005). Knockout and silencing of *SAMT1* in Arabidopsis and *N. benthamiana*, respectively, resulted in leaves with lower levels of αT and total chlorophyll (which also requires SAM for its synthesis) and increased γT content (Bouvier *et al*. 2006), again consistent with reduced import of SAM limiting conversion of γT to αT (and δΤ to βT) by VTE4 (Bouvier *et al*. 2006). Given the strong negative relationship between *samt1* expression and δΤ concentration revealed by TWAS (Supplementary Table S17), we hypothesize that weak expression of *samt1* results in higher accumulation of δΤ by lowering the SAM-dependent activities of VTE4 and/or VTE3 (Figure 1).

The identification of *por1*, *por2*, *vte7*, and *vte5* loci, which predominantly impacted tocopherols, is consistent with them all participating in the generation of PDP for tocopherol synthesis (Albert *et al*. in review; Valentin *et al*. 2006; Diepenbrock *et al*. 2017). The two *por* loci underlie the two largest effect JL-QTL for grain ΣΤ in the maize NAM panel (Diepenbrock *et al*. 2017) and encode a key enzyme for a highly regulated activity in chlorophyll biosynthesis. Diepenbrock *et al*. (2017) found *por1* and *por2* to be correlated expression and effect QTL (ceeQTL) for ΣT; *i.e.*, that the JL allelic effect estimates of these QTL for that trait were significantly associated with expression levels of the respective gene at multiple kernel developmental time points. In concordance with these findings, *por1* and *por2* were also the two most strongly associated genes with ΣΤ via TWAS and FCT in this study. In the Ames panel, an association signal for δΤ localized to *vte7* both by GWAS (5% FDR) and more weakly by TWAS (0.6% of most significant genes). Finally, *vte5*, one of two kinases needed to generate PDP from phytol (Valentin *et al*. 2006; Vom Dorp *et al*. 2015), was associated in GWAS with total tocochromanols (ΣΤΤ3) in maize. In Arabidopsis, *vte5* was one of the strongest associations with seed ΣΤ (Albert *et al*. in review). The association of *por1*, *por2*, *vte7*, and *vte5* with maize grain tocopherol traits in the present study, and VTE7 and VTE5 with seed tocopherol traits in Arabidopsis further strengthens the connection between tocopherol and chlorophyll biosynthesis in seed and suggests it is a major control point for seed tocochromanol variation in both monocots and dicots.

The *dxs1* and *dxs2* genes were only associated in TWAS with tocotrienol, but not tocopherol, traits. Both genes encode the first and committed step of the methylerythritol 4-phosphate (MEP) pathway, suggesting they are key control points in the provision of IPP used to generate GGDP for tocotrienol synthesis. In addition to our finding of an association of *dxs2* transcript abundances with tocotrienol levels, Diepenbrock *et al*. (2017) found *dxs2* to be a ceeQTL for tocotrienol traits (δΤ3, γΤ3, and ΣT3). Our eQTL mapping of *dxs2* expression variation identified a major *trans*-eQTL on chromosome 6 (Supplementary Table S14) that mapped to *phytoene synthase 1* (*psy1*, Zm00001d036345). *psy1* encodes the first and committed step in the biosynthesis of carotenoids (Hirschberg 2001). In Arabidopsis, flux into the carotenoid pathway is controlled by PSY activity via feedback regulation of DXS protein levels (Rodríguez-Villalón *et al*. 2009). Thus, it is possible that *psy1* indirectly affected tocotrienol levels in maize grain, even though we did not find *psy1* to be significantly associated with tocotrienols.

Like *dxs1* and *dxs2*, the three identified genes involved in aromatic head group biosynthesis were specific for tocotrienol traits and include two copies of *arodeH2* (*arodeH2* Zm00001d014734, and *arodeH2* Zm00001d014737) and *hppd1*. ArodeH catalyzes the oxidative decarboxylation of arogenate to tyrosine, which is then transaminated to p-hydroxyphenylpyruvic acid (HPP), the substrate for HPPD. The two *arodeH2* genes are separated by 18 kb and their GWAS associations were with the same peak SNP (5_61159296), located ∼40 kb from Zm00001d014737 and ∼60 kb from Zm00001d014734. Only Zm00001d014737 passed the FCT threshold (< 0.3% vs. > 1.2% for Zm00001d014734), primarily because of its closer proximity to GWAS SNPs with lower *P*-values. A single *cis*-eQTL was declared for Zm00001d014734 (Supplementary Table S15), while Zm00001d014737 lacked significant *cis* or *trans*-eQTL. The current evidence from studies of both the U.S. NAM and Ames panels cannot exclude the possibility that both *arodeH2* genes are involved in head group biosynthesis.

The enzyme encoded by *hppd1* produces HGA, the aromatic headgroup used for synthesis of tocopherols and tocotrienols. In concordance with the findings of Diepenbrock *et al*. (2017), we found *hppd1* to be strongly associated with tocotrienols, but its weak associations with tocopherols in the U.S. NAM panel were not replicated in the Ames panel, likely because the allelic effect sizes for tocopherol traits were below significance in the Ames panel. In agreement with Diepenbrock *et al*. (2017) who found *hppd1* was not a ceeQTL, *hppd1* was also not detected by TWAS in the Ames panel at a 0.5% threshold.

The three identified genes encoding core activities in the tocochromanol pathway (*vte1*, *vte4,* and *hggt1*) impacted tocochromanol phenotypes in a manner consistent with their enzymatic activities (Shintani and DellaPenna 1998; Cahoon *et al*. 2003; Hunter and Cahoon 2007; DellaPenna and Mène-Saffrané 2011). VTE4 methylates γ- and δ-tocopherols to produce α- and β-tocopherols (Shintani and DellaPenna 1998; DellaPenna and Mène-Saffrané 2011). *vte4* was strongly associated with γ and α isoforms in the Ames panel, consistent with prior GWAS reports (Li *et al*. 2012; Lipka *et al*. 2013; Diepenbrock *et al*. 2017; Wang *et al*. 2018; Baseggio *et al*. 2019), and more weakly with δΤ (GWAS and FCT) and ΣΤΤ3 (GWAS).

Similarly, *hggt1* encodes the committed step of tocotrienol biosynthesis and was significantly associated with all tocotrienol traits, reconfirming *hggt1* as the key genetic controller of tocotrienol variation (Lipka *et al*. 2013; Diepenbrock *et al*. 2017; Baseggio *et al*. 2019). However, the relatively weaker associations of *hggt1* with tocopherols detected in the U.S. NAM panel were not reidentified in the Ames panel (Diepenbrock *et al*. 2017). Both *vte4* and *hggt1* were designated as ceeQTL by Diepenbrock *et al*. (2017) and were the highest ranked genes detected in TWAS for αT, αT3, and ΣT3 in the Ames panel, highlighting the importance of regulatory variation in the core tocochromanol pathway. VTE1 converts MPBQ and MGGBQ to δΤ and δΤ3, respectively, and we identified associations of *vte1* with δΤ and δΤ3 at higher resolution (i.e., significant variants closer to the gene) than in other diversity panels (Lipka *et al*. 2013; Baseggio *et al*. 2019) (Table 2 and Supplementary Table S9).

By conducting a statistical analysis that uses a probabilistic method to combine GWAS and eQTL results, we identified a set of 24 unique SNPs at eight candidate causal gene loci that contributed to both GWAS and *cis*-eQTL signals, providing support for an eQTL-mediated mechanism in which variants affect tocochromanol levels. Twenty one of these selected SNPs were located outside (up to 136.5 kb) of the eight genes, with 18 of them upstream of the 5′ end of their respective gene. Even though the selected SNPs are likely not causal themselves and in strong LD with causal variants, these results imply that the underlying causal variants for the colocalized signals predominantly resided in the vicinity of promoters and upstream *cis*-regulatory elements. Although *cis*-eQTL were almost always the largest-effect QTL detected for expression variation of candidate casual genes, our study was limited by statistical power and the multiple testing burden, thus not allowing a more complete genetic dissection of *trans*-eQTL (Albert *et al*. 2018). However, *cis* effects tend to be more stable than *trans* effects across environments, thus *cis*-acting causal variants have the potential to be more transferable across populations when incorporated at the haplotype level in genomic prediction models (Giri *et al*. 2021).

## Conclusions

We identified 13 candidate causal gene loci responsible for controlling natural variation of nine grain tocochromanol phenotypes with encoded activities related to metabolism and metabolite transport. All 13 loci are highly probable to be causal as most have been shown by mutagenesis and transgenic modifications to affect tocochromanol levels in maize and/or other model plant systems. The five novel associations identified for *samt1*, *arodeH2* Zm00001d014737, *dxs1*, *vte5*, and *vte7* together with finer delineation of *vte1*, illustrates the tremendous statistical power and mapping resolution provided by the Ames panel when combining GWAS and TWAS results to study the genetic basis of tocochromanol variation in maize grain. When integrated with the findings of Diepenbrock *et al*. (2017) and Wang *et al*. (2018), there is now a more complete catalog of the key gene targets that connect the tocochromanol and chlorophyll pathways for breeding and engineering of vitamin E and antioxidant levels in maize and other grain crops. The joint statistical analysis of eQTL mapping and GWAS results revealed that *cis*-acting causal variants should be an important consideration when selecting and combining favorable alleles across these key loci to optimize the tocochromanol profile of maize grain for human health and nutrition.

## Data availability

All raw 3′ mRNA-seq data are available from the NCBI Sequence Read Archive under BioProject PRJNA643165. Supplementary Data Sets 1-4 are available on Cyverse (https://de.cyverse.org/data/ds/iplant/home/shared/GoreLab/dataFromPubs/Wu_AmesTocochromonols_2021). Supplementary tables and figures are available at figshare (link). All code is available on Github (https://github.com/GoreLab/Vitamaize_Ames_Toco).

## Acknowledgements

We thank Matt Baseggio, Elise Albert, and other current and past members of the Gore, DellaPenna, Buell, and Yu labs for their efforts in pollination, harvest, and sample preparation.

## Table Captions

**Supplementary Table S1.** Untransformed best linear unbiased estimators (BLUEs) (μg g^−1^) of nine tocochromanol grain phenotypes in the Ames panel

**Supplementary Table S2.** Lambda and constant values used in Box-Cox transformation of nine tocochromanol grain phenotypes in the Ames panel

**Supplementary Table S3.** Transformed best linear unbiased estimators (BLUEs) (μg g^−1^) of nine tocochromanol grain phenotypes in the Ames panel

**Supplementary Table S4.** Summary of samples and genes in the transcriptome-wide association study (TWAS) pipeline

**Supplementary Table S5.** Genomic information for the 126 *a priori* candidate genes in the tocochromanol and chlorophyll biosynthesis pathways. Causal genes that were identified by genetic mapping in this Ames panel are bolded.

**Supplementary Table S6.** Joint-linkage QTL results of nine tocochromanol grain phenotypes analyzed in the maize NAM panel (Diepenbrock *et al*., 2017) uplifted from RefGen_v2 to v4

**Supplementary Table S7.** Genome-wide association study results of nine tocochromanol grain phenotypes analyzed in the maize NAM panel (Diepenbrock *et al*., 2017) uplifted from RefGen_v2 to v4. Only NAM marker variants with resample model inclusion probability (RMIP) ≥ 0.05 are shown and those that reside within joint-linkage QTL support intervals (Supplementary Table S6) are demarcated in the “NAM JL-QTL CSI ID” column.

**Supplementary Table S8.** Rice and Arabidopsis homologs of *samt1*

**Supplementary Table S9.** Genomic information (RefGen_v4) for genes within ± 100 kb of the peak SNPs identified in the genome-wide association study

**Supplementary Table S10.** Genomic information (B73 RefGen_v4) for the top 0.5% of genes identified in the transcriptome-wide association study

**Supplementary Table S11.** Genomic information (RefGen_v4) for the top 0.5% of SNP-gene pairs identified in the Fisher’s combined test

**Supplementary Table S12.** Significant results from the genome-wide association study

**Supplementary Table S13.** Significant results from the genome-wide association study with a multi-locus mixed-model analysis

**Supplementary Table S14.** SnpEff annotation and GERP scores of the 15 GWAS significant SNPs annotated as missense variants

**Supplementary Table S15.** *Cis*- and *trans*-eQTL of the 13 candidate causal genes. Genomic information (RefGen_v4) for the plausible regulating genes within ± 100 kb of the *trans*-eQTL peak SNPs are presented.

**Supplementary Table S16.** All SNPs with CLPP values ≥ 0.01 at candidate causal gene loci

**Supplementary Table S17.** Directionality of association between gene expression (PEER values) and nine tocochromanol phenotypes. Gene-trait pairs that were significant in TWAS (Table 2) are bolded.

**Supplementary Figure S1.** Correlation matrix for untransformed BLUE values for nine tocochromanol grain phenotypes in the Ames panel. Pearson’s correlation coefficients (*r*) calculated with the function ‘*cor*’ in R are presented in the upper triangle, whereas the corresponding *P*-values for the significance of correlations (α = 0.05) are displayed below the diagonal. The untransformed BLUE values were used to represent the true directionality of the relationship between phenotypes.

**Supplementary Figure S2.** Sources of variation for nine tocochromanol grain phenotypes in the Ames panel. The phenotypic variance was statistically partitioned into the following components: G: genotype; G×Y: genotype × year; Y: year; Field: tier(year), pass(tier × year), range(tier × year); Plate: plate(year); and REV: residual error variance. Variance components were estimated by refitting the full model from Equation 1 with genotype as a random effect.

**Supplementary Figure S3.** Genome-wide association study (GWAS), transcriptome-wide association study (TWAS), and Fisher’s combined test (FCT) results for eight tocochromanol phenotypes. A: Upset plot showing the number of overlapping genes between GWAS, TWAS, FCT, and *a priori* pathway genes involved in the biosynthesis of chlorophylls and tocochromanols (Supplementary Table S5). The number of genes located within the U.S. nested association mapping (NAM) joint linkage-quantitative trait loci (JL-QTL) common support interval (CSI) for each phenotype is highlighted in blue in the bar plots. B: Manhattan plots of GWAS, TWAS, and FCT results. Each point represents a SNP or gene with its -log_10_ *P*-value (y-axis) from GWAS, TWAS, and FCT plotted as a function of physical position (Mb, B73 RefGen_v4) across the 10 chromosomes of maize (x-axis). Red horizontal dashed lines indicate the thresholds of top 0.02%, top 0.5% and top 0.5% for GWAS, TWAS, and FCT, respectively. Causal genes (Table 2) that are within 100 kb of a top 0.02% GWAS peak SNP or ranked top 0.5% in TWAS or FCT are highlighted with red dots and labeled in black in the Manhattan plots. Causal genes that are within 1 Mb of a top 0.02% GWAS peak SNP are labeled in gray. Novel associations are marked with a solid line, black rectangle.

**Supplementary Figure S4.** Manhattan plots of expression quantitative trait loci (eQTL) mapping results of 12 candidate causal genes. Each point represents a SNP with its -log_10_ *P*-value (y-axis) from a mixed linear model analysis plotted as a function of physical position (B73 RefGen_v4) across the 10 chromosomes of maize (x-axis). The red horizontal dashed line indicates the significant threshold after Bonferroni correction (α = 0.05). The green vertical line indicates the genomic position of each candidate causal gene.

**Supplementary Figure S5.** Genome-wide association study (GWAS), expression quantitative trait loci (eQTL), and eQTL and GWAS CAusal Variants Identification in Associated Regions (eCAVIAR) results of 22 gene-phenotype pairs. A, Three local Manhattan plots (± 100 kb) showing the GWAS, eQTL, and eCAVIAR results of each gene-phenotype pair. Each point represents a SNP with its -log_10_ *P*-value (y-axis) from a mixed linear model analysis in GWAS or eQTL, or the CLPP value from eCAVIAR, plotted as a function of physical position (Mb, B73 RefGen_v4) on x-axis. The blue rectangle represents the physical position of the gene. The red dot represents the SNP with the highest CLPP value, and the orange dot represents the other SNPs with a CLPP value passing the ≥ 0.01 threshold. B, Pairwise linkage disequilibrium (LD, *r^2^*) of all SNPs included in A, with the LD between SNPs with CLPP values ≥ 0.01 indicated in the bottom left.

## Supplementary Data Sets captions

**Supplementary Data Set 1.** This folder contains the marker genotype and kinship matrix used for genome-wide association studies

**Supplementary Data Set 2.** This folder contains the marker genotype and sample call rate results used for line selection of the 2018 field experiment for collecting developing grain samples for transcriptomic profiling

**Supplementary Data Set 3.** This folder contains the probabilistic estimation of expression residual (PEER) values used for transcriptome-wide association studies and expression quantitative trait loci mapping

**Supplementary Data Set 4.** This folder contains the marker genotype matrix used for expression quantitative trait loci mapping and the kinship matrix used for both transcriptome-wide association studies and expression quantitative trait loci mapping

## Supplementary Methods

### Statistical modeling and filtering of phenotypic data

To screen the raw high-performance liquid chromatography (HPLC) data for significant outliers, we fitted a mixed linear model for each tocochromanol phenotype in ASReml-R version 3.0 (Gilmour *et al*. 2009). The full model (Equation 1) fitted to the data was as follows:

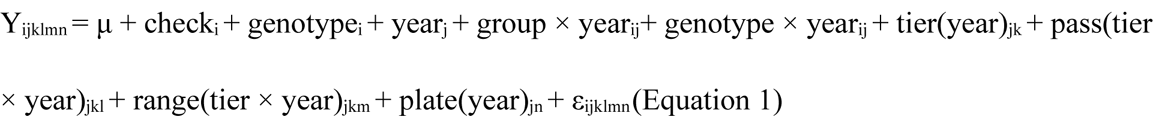

in which Y_ijklmn_ is an individual phenotypic observation; μ is the grand mean; check_i_ is the fixed effect for the B73 check, where it is set to 0 if the genotype is a non-check line; genotype_i_ is the fixed effect of the ith genotype (non-check line), where it is set as 0 and omitted if the observation is of the B73 check; year_j_ is the effect of the jth year; group × year_ij_ is the interaction term between the ith group and jth year, where group_i_ is an indicator variable with two levels that indicates whether the observation is of a B73 check or non-check line; genotype × year_ij_ is the effect of the interaction between the ith genotype (non-check line) and jth year, which is not included in the model for the B73 check; tier(year)_jk_ is the effect of the kth tier within the jth year; pass(tier × year)_jkl_ is the effect of the lth pass within the kth tier within the jth year; range(tier × year)_jkm_ is the effect of the mth range within the kth tier within the jth year; plate(year)_jn_ is the effect of the nth HPLC autosampler plate within the jth year; and ε_ijklmn_ is the residual error effect assumed to be independently and identically distributed (i.i.d.) according to a normal distribution with mean zero and variance σ_ε_ , that is ∼iid N(0, σ_ε_ ). Of these terms, μ, check, and genotype were modeled as fixed effects, while all other terms were modeled as random effects. Studentized deleted residuals (Neter *et al*. 1996) generated by the model were used to remove 147 significant outliers (where an outlier is a single plot observation for a single trait) at a Bonferroni adjusted significance threshold of α = 0.05. With the outlier-screened raw HPLC data set, we generated best linear unbiased estimator (BLUE) values for 1,762 inbred lines across years (Supplementary Table S1) by fitting the full model (Equation 1) in ASReml-R version 3.0 (Gilmour *et al*. 2009). The full model was refitted with genotype as a random effect to generate variance component estimates for the calculation of heritability on a line-mean basis (Holland *et al*. 2003; Hung *et al*. 2012).

### Genotype data processing and imputation

The marker genotype imputation approach implemented in Wu *et al*. (2021) was used to generate a high-density single-nucleotide polymorphism (SNP) marker set in B73 RefGen_v4 coordinates for the Ames panel. To construct the target SNP genotype set, unimputed genotyping-by-sequencing (GBS) SNP genotypes scored at 943,455 loci in the Ames panel by Romay *et al*. (2013) were downloaded from CyVerse (ZeaGBSv27_publicSamples_raw_AGPv4-181023.vcf.gz, available at http://datacommons.cyverse.org/browse/iplant/home/shared/panzea/genotypes/GBS/v27), which provided 1,779 GBS samples for 1,493 of the 1,497 lines that had best linear unbiased estimator (BLUE) values for tocochromanol phenotypes. Given that there were 220 lines with more than one corresponding GBS sample having a call rate ≥ 20%, we followed the approach of Wu *et al*. (2021) to merge two or more GBS samples from the same line. Briefly, a stringently filtered SNP set (call rate ≥ 50%, % heterozygosity ≤ 10%, index of panmixia FIT ≥ 0.8, minor allele frequency ≥ 0.01 and linkage disequilibrium *r^2^*≤ 0.2) of 32,267 SNPs derived from the Romay *et al*. (2013) unimputed marker data set was used to calculate average pairwise identity-by-state (IBS) between multiple samples of the same line using PLINK version 1.9 (Purcell *et al*. 2007). A total of 19 lines with a mean IBS value < 0.95 for all within-line sample comparisons were removed from the analysis, followed by consensus genotype calling for the remaining 201 lines. Collectively, the final target data set consisted of 443,419 biallelic GBS SNPs scored on a retained 1,462 lines with a call rate ≥ 0.2%, heterozygosity ≤ 10%, and inbreeding coefficient (F) ≥ 0.8. All heterozygous genotype calls were set to missing prior to imputation.

The reference SNP genotype set, which was identical to that constructed in Wu *et al*. (2021), consisted of 14,613,169 SNPs derived from maize HapMap 3.2.1 (Bukowski *et al*. 2018). In BEAGLE v5.0 (Browning *et al*. 2018) with parameters as previously specified in Wu *et al*. (2021), the genotypes at the 14,613,169 SNP loci were imputed based on 443,419 GBS SNPs (target set) in the 1,462 Ames panel lines. This data set of 14,613,169 loci served as the foundation for the subsetting of markers for all of the quantitative genetic analyses conducted in this study.

### Expression data set quality control

To verify the quality and integrity of the samples, SNPs were called using the 3′ QuantSeq read alignments and compared to SNP calls from a 942 maize line RNA-Seq data set (WiDiv-942 panel) (Gage *et al*. 2019). In total, 375 lines overlapped with the WiDiv-942 panel, for a total of 430 3′ QuantSeq samples and 54 positive controls. First, 3′ QuantSeq reads were mapped to the B73 RefGen_v4 assembly (Jiao *et al*. 2017) following the HISAT2 mapping protocol indicated above. Duplicate reads were identified and marked using Picard tools MarkDuplicates version 2.20.8 (https://broadinstitute.github.io/picard/). Output was sorted using SAMTools sort version 1.9 and a pileup file created using SAMTools mpileup with BAQ computation disabled (-B) and alignments with a mapQ less than 60 were omitted (-q 60), allowing for only unique alignments to be processed. Only positions with a base quality of greater than or equal to 20 were included and all insertions and deletions were discarded. Genotype calls were made at a position in an individual if the coverage was at least five reads, but not greater than 500 reads, and the allele made up greater than 3% of the calls at that position in the individual. If more than two alleles passed the coverage and frequency cutoff, the position was scored as heterozygous and set to missing data only when calculating percent identity. After removing positions from the WiDiv-942 SNP matrix that were not called in the 3′ QuantSeq data set, there were 919,074 remaining positions. Percent identity between the same line in the two data sets was calculated by taking the number of positions that had the same genotype call at a position divided by the total number of positions excluding missing data positions in either data set.

Stringent filtering was employed to curate the final expression data set and ensure it contained high quality data. Samples were filtered out based on the following criteria: sampling concerns such as moldy kernels etc. (12 samples removed), number of cleaned reads were below 5 million (1 sample removed), a HISAT2 alignment rate of less than or equal to 65% (17 samples removed), a Pearson’s correlation value (*r*) less than 0.90 with 40 or more samples (3 samples removed), samples that had less than 95% identity when compared to their high confidence WiDiv-942 panel counterpart during genotype confirmation assessment (15 samples removed), and finally removal of samples that had an heterozygosity greater than or equal to 10% (339 samples removed). This final heterozygosity filter was employed to remove samples that were contaminated by spillover during library construction at the Cornell Institute of Biotechnology’s Genomics Facility. This stringent heterozygosity filtering was employed to ensure the final data set was free of contaminating reads that may have impacted downstream analysis. The final data set of 784 high confidence, high quality samples included 43 B73 positive control samples and 741 collected field samples of check and noncheck lines. The B73 positive controls were used during data processing for quality control. Only the 741 check and noncheck samples were used for downstream expression data analysis.

### Statistical modeling of expression data

To account for the potential effect of different amounts of accumulated heat units on kernel development, growing degree days (GDD, Bollero *et al*. 1996) from pollination to fresh-harvest at the ∼23 DAP time point for each ear was included as a model term when calculating BLUE expression values.

For each gene, BLUE values were generated for each of the 664 non-check lines in ASReml-R version 3.0 (Gilmour *et al*. 2009) as follows:

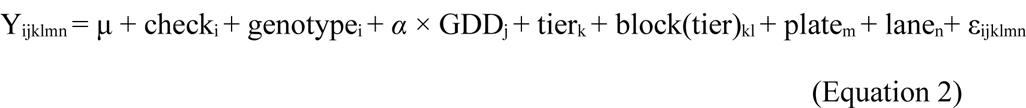

in which Y_ijklmn_ is an individual *rlog*-transformed value; μ is the grand mean; check_i_ is the fixed effect for the ith check, where it is set to 0 if the genotype is a non-check line; genotype_i_ is the fixed effect of the ith genotype (non-check line), where it is set as 0 and omitted if the ith observation is of a check line; *α* is a scalar regression coefficient for the GDD value of ears harvested on the jth day (GDD_j_); tier_k_ is the kth tier; block(tier)_kl_ is the lth block in the kth tier; plate_m_ is the mth RNA sample plate; lane_n_ is the nth lane on an Illumina sequencer (minimum unit of the RNAseq run); and ε_ijklmn_ is the residual error effect assumed to be ∼iid N(0, σ_ε_ ). With the exception of the grand mean, check, genotype and GDD, all terms were fitted as random effects.

### Expression data analysis for *vte7*

Given that the *vte7* locus consists of tandemly duplicated genes (Zm00001d006778 and Zm00001d006779) with high pairwise nucleotide sequence identity (> 99%) in the B73 RefGen_v4 assembly (Jiao *et al*. 2017), the reads pertaining to these two genes were not uniquely mappable with our standard expression abundance determination bioinformatic pipeline. Therefore, to calculate the transcript abundances at the *vte7* locus, the number of read alignments to the two gene models using multi-mapping reads were summed and normalized to counts per million alignments (CPMA) as (total count within both loci/total reads aligned)*1,000,000. Next, the CPMA values were fitted with the Equation 2 model to generate BLUE values, which were then screened for outliers with Studentized deleted residuals (Neter *et al*. 1996) to produce the final *vte7* data set (Supplementary Data Set 3).

## Literature cited

Albert E., Kim S Deason, Y. Bao, M. Magallanes-Lundback, B. Danilo, et al., Genome-wide association identifies a missing hydrolase for tocopherol biosynthesis in plants. In review.

Albert F. W., J. S. Bloom, J. Siegel, L. Day, and L. Kruglyak, 2018 Genetics of trans-regulatory variation in gene expression. Elife 7: e35471.

Anders S., P. T. Pyl, and W. Huber, 2015 HTSeq—a Python framework to work with high-throughput sequencing data. Bioinformatics 31: 166–169.

Baseggio M., M. Murray, M. Magallanes-Lundback, N. Kaczmar, J. Chamness, et al., 2019 Genome-wide association and genomic prediction models of tocochromanols in fresh sweet corn kernels. The Plant Genome 12. https://doi.org/10.3835/plantgenome2018.06.0038

Benjamini Y., and Y. Hochberg, 1995 Controlling the false discovery rate: A practical and powerful approach to multiple testing. J. R. Stat. Soc. Series B Stat. Methodol. 57: 289–300.

Bouvier F., A. Rahier, and B. Camara, 2005 Biogenesis, molecular regulation and function of plant isoprenoids. Prog. Lipid Res. 44: 357–429.

Bouvier F., N. Linka, J.-C. Isner, J. Mutterer, A. P. M. Weber, et al., 2006 Arabidopsis SAMT1 defines a plastid transporter regulating plastid biogenesis and plant development. Plant Cell 18: 3088–3105.

Box G. E. P., and D. R. Cox, 1964 An analysis of transformations. J. R. Stat. Soc. Series B Stat. Methodol. 26: 211–243.

Browning B. L., Y. Zhou, and S. R. Browning, 2018 A one-penny imputed genome from next-generation reference panels. Am. J. Hum. Genet. 103: 338–348.

Bukowski R., X. Guo, Y. Lu, C. Zou, B. He, et al., 2018 Construction of the third-generation *Zea mays* haplotype map. Gigascience 7: 1–12.

Cahoon E. B., S. E. Hall, K. G. Ripp, T. S. Ganzke, W. D. Hitz, et al., 2003 Metabolic redesign of vitamin E biosynthesis in plants for tocotrienol production and increased antioxidant content. Nat. Biotechnol. 21: 1082–1087.

Cheng Z., S. Sattler, H. Maeda, Y. Sakuragi, D. A. Bryant, et al., 2003 Highly divergent methyltransferases catalyze a conserved reaction in tocopherol and plastoquinone synthesis in cyanobacteria and photosynthetic eukaryotes. Plant Cell 15: 2343–2356.

Cingolani P., A. Platts, L. L. Wang, M. Coon, T. Nguyen, et al., 2012 A program for annotating and predicting the effects of single nucleotide polymorphisms, SnpEff: SNPs in the genome of Drosophila melanogaster strain w1118; iso-2; iso-3. Fly 6: 80–92.

Davydov E. V., D. L. Goode, M. Sirota, G. M. Cooper, A. Sidow, et al., 2010 Identifying a high fraction of the human genome to be under selective constraint using GERP++. PLoS Comput. Biol. 6: e1001025.

DellaPenna D., 2005 A decade of progress in understanding vitamin E synthesis in plants. Plant Physiol. 162: 729–737.

DellaPenna D., and L. Mène-Saffrané, 2011 Vitamin E, pp. 179–227 in Advances in Botanical Research, edited by Rébeillé F., Douce R. Elsevier Ltd., Amsterdam, The Netherlands.

Dewey M., 2019 metap: Meta-analysis of significance values. R package version 1.1

Diepenbrock C. H., C. B. Kandianis, A. E. Lipka, M. Magallanes-Lundback, B. Vaillancourt, et al., 2017 Novel loci underlie natural variation in vitamin E levels in maize grain. Plant Cell 29: 2374–2392.

Diepenbrock C. H., D. C. Ilut, M. Magallanes-Lundback, C. B. Kandianis, A. E. Lipka, et al., 2021 Eleven biosynthetic genes explain the majority of natural variation in carotenoid levels in maize grain. Plant Cell 33: 882–900.

Endelman J. B., 2011 Ridge regression and other kernels for genomic selection with R package rrBLUP. The Plant Genome 4: 250.

Ferguson J. N., S. B. Fernandes, B. Monier, N. D. Miller, D. Allen, et al., 2021 Machine learning-enabled phenotyping for GWAS and TWAS of WUE traits in 869 field-grown sorghum accessions. Plant Physiol. 187: 1481–1500.

Ford E. S., R. L. Schleicher, A. H. Mokdad, U. A. Ajani, and S. Liu, 2006 Distribution of serum concentrations of α-tocopherol and γ-tocopherol in the US population. Am. J. Clin. Nutr. 84: 375–383.

Giri A., M. Khaipho-Burch, E. S. Buckler, and G. P. Ramstein, 2021 Haplotype associated RNA expression (HARE) improves prediction of complex traits in maize. PLoS Genet. 17: e1009568.

Hershberger J., R. Tanaka, J. C. Wood, N. Kaczmar, D. Wu, et al., 2022 Transcriptome-wide association and prediction for carotenoids and tocochromanols in fresh sweet corn kernels. The Plant Genome e20197. https://doi.org/10.1002/tpg2.20197

Hirsch C. N., J. M. Foerster, J. M. Johnson, R. S. Sekhon, G. Muttoni, et al., 2014 Insights into the maize pan-genome and pan-transcriptome. Plant Cell 26: 121–135.

Hirschberg J., 2001 Carotenoid biosynthesis in flowering plants. Curr. Opin. Plant Biol. 4: 210– 218.

Hormozdiari F., M. van de Bunt, A. V. Segrè, X. Li, J. W. J. Joo, et al., 2016 Colocalization of GWAS and eQTL signals detects target genes. Am. J. Hum. Genet. 99: 1245–1260.

Hunter S. C., and E. B. Cahoon, 2007 Enhancing vitamin E in oilseeds: unraveling tocopherol and tocotrienol biosynthesis. Lipids 42: 97–108.

Jiao Y., P. Peluso, J. Shi, T. Liang, M. C. Stitzer, et al., 2017 Improved maize reference genome with single-molecule technologies. Nature 546: 524–527.

Kim D., J. M. Paggi, C. Park, C. Bennett, and S. L. Salzberg, 2019 Graph-based genome alignment and genotyping with HISAT2 and HISAT-genotype. Nat. Biotechnol. 37: 907– 915.

Kistler L., S. Y. Maezumi, J. Gregorio de Souza, N. A. S. Przelomska, F. Malaquias Costa, et al., 2018 Multiproxy evidence highlights a complex evolutionary legacy of maize in South America. Science 362: 1309–1313.

Knekt P., A. Reunanen, R. Järvinen, R. Seppänen, M. Heliövaara, et al., 1994 Antioxidant vitamin intake and coronary mortality in a longitudinal population study. Am. J. Epidemiol. 139: 1180–1189.

Kremling K. A. G., C. H. Diepenbrock, M. A. Gore, E. S. Buckler, and N. B. Bandillo, 2019 Transcriptome-wide association supplements genome-wide association in *Zea mays*. G3 9: 3023–3033.

Kurtz S., 2010 The Vmatch large scale sequence analysis software - a manual. http://www.vmatch.de/virtman.pdf

Kushi L. H., A. R. Folsom, R. J. Prineas, P. J. Mink, Y. Wu, et al., 1996 Dietary antioxidant vitamins and death from coronary heart disease in postmenopausal women. N. Engl. J. Med. 334: 1156–1162.

Li H., B. Handsaker, A. Wysoker, T. Fennell, J. Ruan, et al., 2009 The sequence alignment/map format and SAMtools. Bioinformatics 25: 2078–2079.

Li Q., X. Yang, S. Xu, Y. Cai, D. Zhang, et al., 2012 Genome-wide association studies identified three independent polymorphisms associated with α-tocopherol content in maize kernels. PLoS One 7: e36807.

Li H., Z. Peng, X. Yang, W. Wang, J. Fu, et al., 2013 Genome-wide association study dissects the genetic architecture of oil biosynthesis in maize kernels. Nat. Genet. 45: 43–50.

Li D., Q. Liu, and P. S. Schnable, 2021 TWAS results are complementary to and less affected by linkage disequilibrium than GWAS. Plant Physiol. 186: 1800–1811.

Lin H.-Y., Q. Liu, X. Li, J. Yang, S. Liu, et al., 2017 Substantial contribution of genetic variation in the expression of transcription factors to phenotypic variation revealed by eRD-GWAS. Genome Biol. 18: 192.

Lipka A. E., F. Tian, Q. Wang, J. Peiffer, M. Li, et al., 2012 GAPIT: genome association and prediction integrated tool. Bioinformatics 28: 2397–2399.

Lipka A. E., M. A. Gore, M. Magallanes-Lundback, A. Mesberg, H. Lin, et al., 2013 Genome-wide association study and pathway-level analysis of tocochromanol levels in maize grain. G3 3: 1287–1299.

Love M. I., W. Huber, and S. Anders, 2014 Moderated estimation of fold change and dispersion for RNA-seq data with DESeq2. Genome Biol. 15: 550.

Lozano R., E. Gazave, J. P. R. dos Santos, M. G. Stetter, R. Valluru, et al., 2021 Comparative evolutionary genetics of deleterious load in sorghum and maize. Nature Plants 7: 17–24.

Majewski J., and T. Pastinen, 2011 The study of eQTL variations by RNA-seq: from SNPs to phenotypes. Trends Genet. 27: 72–79.

Martin M., 2011 Cutadapt removes adapter sequences from high-throughput sequencing reads. EMBnet.journal 17: 10–12.

McBurney M. I., E. A. Yu, E. D. Ciappio, J. K. Bird, M. Eggersdorfer, et al., 2015 Suboptimal serum α-tocopherol concentrations observed among younger adults and those depending exclusively upon food sources, NHANES 2003–2006. PLoS One 10: e0135510.

McMullen M. D., S. Kresovich, H. S. Villeda, P. Bradbury, H. Li, et al., 2009 Genetic properties of the maize nested association mapping population. Science 325: 737–740.

Neter J., M. H. Kutner, C. J. Nachtsheim, and W. Wasserman, 1996 Applied linear statistical models. Irwin Chicago.

Palmieri L., R. Arrigoni, E. Blanco, F. Carrari, M. I. Zanor, et al., 2006 Molecular identification of an arabidopsis S-adenosylmethionine transporter: Analysis of organ distribution, bacterial expression, reconstitution into liposomes, and functional characterization. Plant Physiol. 142: 855–865.

Pasaniuc B., and A. L. Price, 2017 Dissecting the genetics of complex traits using summary association statistics. Nat. Rev. Genet. 18: 117–127.

Pignon C. P., S. B. Fernandes, R. Valluru, N. Bandillo, R. Lozano, et al., 2021 Phenotyping stomatal closure by thermal imaging for GWAS and TWAS of water use efficiency-related genes. Plant Physiol. 187: 2544–2562.

Porfirova S., E. Bergmuller, S. Tropf, R. Lemke, and P. Dormann, 2002 Isolation of an *Arabidopsis* mutant lacking vitamin E and identification of a cyclase essential for all tocopherol biosynthesis. Proc. Natl. Acad. Sci. U. S. A. 99: 12495–12500.

Purcell S., B. Neale, K. Todd-Brown, L. Thomas, M. A. R. Ferreira, et al., 2007 PLINK: A tool set for whole-genome association and population-based linkage analyses. Am. J. Hum. Genet. 81: 559–575.

Ramstein G. P., S. J. Larsson, J. P. Cook, J. W. Edwards, E. S. Ersoz, et al., 2020 Dominance effects and functional enrichments improve prediction of agronomic traits in hybrid maize. Genetics 215: 215–230.

R Core Team, 2018 An Introduction to R

Rodgers-Melnick E., P. J. Bradbury, R. J. Elshire, J. C. Glaubitz, C. B. Acharya, et al., 2015 Recombination in diverse maize is stable, predictable, and associated with genetic load. Proc. Natl. Acad. Sci. U. S. A. 112: 3823–3828.

Rodríguez-Villalón A., E. Gas, and M. Rodríguez-Concepción, 2009 Phytoene synthase activity controls the biosynthesis of carotenoids and the supply of their metabolic precursors in dark-grown Arabidopsis seedlings. Plant J. 60: 424–435.

Romay M. C., M. J. Millard, J. C. Glaubitz, J. A. Peiffer, K. L. Swarts, et al., 2013 Comprehensive genotyping of the USA national maize inbred seed bank. Genome Biol. 14: R55.

Sattler S. E., L. U. Gilliland, M. Magallanes-Lundback, M. Pollard, and D. DellaPenna, 2004 Vitamin E is essential for seed longevity and for preventing lipid peroxidation during germination. Plant Cell 16: 1419–1432.

Schwarz G., 1978 Estimating the dimension of a model. The Annals of Statistics 6: 461–464.

Segura V., B. J. Vilhjálmsson, A. Platt, A. Korte, Ü. Seren, et al., 2012 An efficient multi-locus mixed-model approach for genome-wide association studies in structured populations. Nat. Genet. 44: 825–830.

Sen C. K., S. Khanna, and S. Roy, 2006 Tocotrienols: Vitamin E beyond tocopherols. Life Sci. 78: 2088–2098.

Shintani D., and D. DellaPenna, 1998 Elevating the vitamin E content of plants through metabolic engineering. Science 282: 2098–2100.

Stegle O., L. Parts, M. Piipari, J. Winn, and R. Durbin, 2012 Using probabilistic estimation of expression residuals (PEER) to obtain increased power and interpretability of gene expression analyses. Nat. Protoc. 7: 500–507.

Sun G., C. Zhu, M. H. Kramer, S.-S. Yang, W. Song, et al., 2010 Variation explained in mixed-model association mapping. Heredity 105: 333–340.

Traber M. G., 2012 Vitamin E, in Present Knowledge in Nutrition, edited by Erdman J. W. Jr, MacDonald I. A., Zeisel S. H. John Wiley & Sons.

Valentin H. E., K. Lincoln, F. Moshiri, P. K. Jensen, Q. Qi, et al., 2006 The *Arabidopsis* vitamin E pathway *gene5-1* mutant reveals a critical role for phytol kinase in seed tocopherol biosynthesis. Plant Cell 18: 212–224.

Van Eenennaam A. L., K. Lincoln, T. P. Durrett, H. E. Valentin, C. K. Shewmaker, et al., 2003 Engineering vitamin E content: From Arabidopsis mutant to soy oil. Plant Cell 15: 3007– 3019.

VanRaden P. M., 2008 Efficient methods to compute genomic predictions. J. Dairy Sci. 91: 4414–4423.

Vom Dorp K., G. Hölzl, C. Plohmann, M. Eisenhut, M. Abraham, et al., 2015 Remobilization of phytol from chlorophyll degradation is essential for tocopherol synthesis and growth of Arabidopsis. Plant Cell 27: 2846–2859.

Wan C. Y., and T. A. Wilkins, 1994 A modified hot borate method significantly enhances the yield of high-quality RNA from cotton (*Gossypium hirsutum* L.). Anal. Biochem. 223: 7–12.

Wang H., S. Xu, Y. Fan, N. Liu, W. Zhan, et al., 2018 Beyond pathways: genetic dissection of tocopherol content in maize kernels by combining linkage and association analyses. Plant Biotechnol. J. 16: 1464–1475.

Wu D., R. Tanaka, X. Li, G. P. Ramstein, S. Cu, et al., 2021 High-resolution genome-wide association study pinpoints metal transporter and chelator genes involved in the genetic control of element levels in maize grain. G3 11. https://doi.org/10.1093/g3journal/jkab059

Yang J., S. Mezmouk, A. Baumgarten, E. S. Buckler, K. E. Guill, et al., 2017 Incomplete dominance of deleterious alleles contributes substantially to trait variation and heterosis in maize. PLoS Genet. 13: e1007019.

Yu J., G. Pressoir, W. H. Briggs, I. Vroh Bi, M. Yamasaki, et al., 2006 A unified mixed-model method for association mapping that accounts for multiple levels of relatedness. Nat. Genet. 38: 203–208.

Yu J., J. B. Holland, M. D. McMullen, and E. S. Buckler, 2008 Genetic design and statistical power of nested association mapping in maize. Genetics 178: 539–551.

Zhan W., J. Liu, Q. Pan, H. Wang, S. Yan, et al., 2019 An allele of *ZmPORB2* encoding a protochlorophyllide oxidoreductase promotes tocopherol accumulation in both leaves and kernels of maize. Plant J. 100: 114–127.

Zhang Z., E. Ersoz, C.-Q. Lai, R. J. Todhunter, H. K. Tiwari, et al., 2010 Mixed linear model approach adapted for genome-wide association studies. Nat. Genet. 42: 355–360.

## Literature cited

Bollero G. A., D. G. Bullock, and S. E. Hollinger, 1996 Soil temperature and planting date effects on corn yield, leaf area, and plant development. Agron. J. 88: 385–390.

Gage J. L., B. Vaillancourt, J. P. Hamilton, N. C. Manrique-Carpintero, T. J. Gustafson, et al., 2019 Multiple maize reference genomes impact the identification of variants by genome-wide association study in a diverse inbred panel. Plant Genome 12. https://doi.org/10.3835/plantgenome2018.09.0069

Gilmour A. R., B. J. Gogel, B. R. Cullis, R. Thompson, D. Butler, et al., 2009 ASReml user guide release 3.0. VSN International Ltd, Hemel Hempstead, UK.

Holland J. B., W. E. Nyquist, and C. T. Cervantes-Martínez, 2003 Estimating and interpreting heritability for plant breeding: an update. Plant Breed. Rev. 22: 9–112.

Hung H.-Y., C. Browne, K. Guill, N. Coles, M. Eller, et al., 2012 The relationship between parental genetic or phenotypic divergence and progeny variation in the maize nested association mapping population. Heredity 108: 490–499.

Wu D., R. Tanaka, X. Li, G. P. Ramstein, S. Cu, et al., 2021 High-resolution genome-wide association study pinpoints metal transporter and chelator genes involved in the genetic control of element levels in maize grain. G3. https://doi.org/10.1093/g3journal/jkab059

